# Interferon receptor gene dosage determines diverse hallmarks of Down syndrome

**DOI:** 10.1101/2022.02.03.478982

**Authors:** Katherine A. Waugh, Ross Minter, Jessica Baxter, Congwu Chi, Kathryn D. Tuttle, Neetha P. Eduthan, Matthew D. Galbraith, Kohl T. Kinning, Zdenek Andrysik, Paula Araya, Hannah Dougherty, Lauren N. Dunn, Michael Ludwig, Kyndal A. Schade, Dayna Tracy, Keith P. Smith, Ross E. Granrath, Nicolas Busquet, Santosh Khanal, Ryan D. Anderson, Liza L. Cox, Belinda Enriquez Estrada, Angela L. Rachubinski, Hannah R. Lyford, Eleanor C. Britton, David J. Orlicky, Jennifer L. Matsuda, Kunhua Song, Timothy C. Cox, Kelly D. Sullivan, Joaquin M. Espinosa

**Affiliations:** Linda Crnic Institute for Down Syndrome, University of Colorado Anschutz Medical Campus; Aurora, CO, USA; Department of Pharmacology, University of Colorado Anschutz Medical Campus; Aurora, CO, USA; Division of Cardiology, Department of Medicine, University of Colorado Anschutz Medical Campus; Aurora, CO, USA; Gates Center for Regenerative Medicine and Stem Cell Biology, University of Colorado Anschutz Medical Campus; Aurora, CO, USA; The Consortium for Fibrosis Research & Translation, University of Colorado Anschutz Medical Campus; Aurora, CO, USA; Animal Behavior Core, NeuroTechnology Center, University of Colorado Anschutz Medical Campus; Aurora, CO, USA., University of Colorado Anschutz Medical Campus; Aurora, CO, USA; Department of Neurology, University of Colorado Anschutz Medical Campus; Aurora, CO, USA; Department of Oral and Craniofacial Sciences, University of Missouri-Kansas City; Kansas City, MO, USA; Department of Pediatrics, University of Missouri-Kansas City; Kansas City, MO, USA; Department of Pathology, University of Colorado Anschutz Medical Campus; Aurora, CO, USA; Department of Immunology and Genomic Medicine, National Jewish Health; Denver, CO, USA; Department of Pediatrics, Section of Developmental Biology, University of Colorado Anschutz Medical Campus; Aurora, CO, USA

**Keywords:** Interferon, trisomy 21, Down syndrome, CRISPR/Cas9, genome editing, inflammation, heart defects, craniofacial morphology, developmental delays, cognition

## Abstract

Trisomy 21 causes Down syndrome, a condition characterized by cognitive impairments, immune dysregulation, and atypical morphogenesis. Using whole blood transcriptome analysis, we demonstrate that specific overexpression of four interferon receptors encoded on chromosome 21 associates with chronic interferon hyperactivity and systemic inflammation in Down syndrome. To define the contribution of interferon receptor overexpression to Down syndrome phenotypes, we used genome editing to correct interferon receptor gene dosage in mice carrying triplication of a large genomic region orthologous to human chromosome 21. Normalization of interferon receptor copy number attenuated lethal antiviral responses, prevented heart malformations, decreased developmental delays, improved cognition and normalized craniofacial anomalies. Therefore, interferon receptor gene dosage determines major hallmarks of Down syndrome, indicating that trisomy 21 elicits an interferonopathy amenable to therapeutic intervention.

**One-Sentence Summary:** Correction of interferon receptor gene dosage rescues multiple key phenotypes in a mouse model of trisomy 21.

## Main Text

Triplication of human chromosome 21 (trisomy 21, T21) occurs 1 in ∼700 live births, causing Down syndrome (DS) (*1, 2*). People born with DS experience developmental delays, cognitive impairment, craniofacial abnormalities, and high rates of congenital heart defects (*3*). These individuals also display a distinct clinical profile, exhibiting lower risk of most solid malignancies, atherosclerosis, and hypertension, along with increased risk of severe respiratory viral infections, Alzheimer’s disease, leukemia, autism spectrum disorders, and diverse autoimmune diseases (*3–5*). Therefore, research on the mechanisms driving these phenomena will benefit not only people with DS but also those in the general population affected by conditions modulated by T21.

Interferon (IFN) signaling, a key component of the innate immune system, is hyperactive in people with DS (*6*). Upon receptor binding, IFN ligands induce JAK/STAT signaling and a downstream transcriptional program to elicit diverse context-dependent cellular responses, including restriction of viral replication, decreased cell proliferation, apoptosis, metabolic reprogramming, and immune activation (*7*). The mechanisms driving IFN hyperactivity in DS are debated. Notably, four of the six IFN receptors (*IFNRs*) are encoded on human chromosome 21 (HSA21) and overexpressed in diverse cell types with T21: *IFNAR1* and *IFNAR2* for Type I IFNs, *IFNGR2* for Type II IFNs, and *IL10RB* for Type III IFNs (*6, 8*). Cells with T21 are hypersensitive to IFN stimulation (*6, 9–11*) and this hypersensitivity is rescued *in vitro* by reducing *IFNR* copy number (*10*), indicating that *IFNR* triplication could drive elevated IFN responses in DS. However, multiple constitutive trisomies, including T21, have been shown to elevate IFN signaling through accumulation of cytosolic dsDNA and activation of the cGAS-STING pathway (*12*). The consequences of chronic IFN hyperactivity in DS remain to be elucidated. Importantly, mutations leading to chronic elevation in IFN signaling cause interferonopathies, a group of monogenic disorders that share some developmental and clinical features with DS (*13, 14*). Therefore, elucidating the exact mechanism driving IFN hyperactivity in DS and its contribution to DS phenotypes could identify targeted therapeutic strategies to counteract the detrimental effects of T21.

In this study, we utilized whole blood transcriptome analysis in a large cohort of individuals with DS to define associations between overexpression of individual HSA21 genes and inflammatory markers in DS, which revealed that only a select subset of triplicated genes, including the four *IFNRs,* associate with IFN hyperactivity and inflammation. We then employed genome editing to normalize gene dosage for all four *Ifnrs* in a mouse model of DS, which revealed that *Ifnr* copy number determines multiple phenotypes of DS. These results identify aberrant IFN signaling as a common node underlying the etiology of vastly diverse hallmarks of the most common human chromosomal abnormality, with clear therapeutic implications for the clinical management of DS, while also advancing the understanding of the impacts of dysregulated cytokine signaling in human development.

### Inflammatory signatures associate with overexpression of specific triplicated genes in DS

In order to investigate mechanisms by which T21 causes IFN hyperactivity in DS, we analyzed the datasets generated by the Crnic Institute Human Trisome Project (HTP), which include whole blood transcriptome and matched plasma immune markers in 304 individuals with T21 versus 96 typical controls (D21) (see **Methods**, **Fig. S1A, Data S1**). These datasets, which were generated from biospecimens collected from individuals ages 1-61 (**Fig. S1B**), enabled us to complete a correlation study between overexpression of individual genes encoded on HSA21 and markers of immune dysregulation in both the transcriptome and proteome across the lifespan. The transcriptome analysis detected 173 mRNAs and lncRNAs encoded on HSA21, 90% of which were significantly upregulated in T21 relative to controls, with a mean fold change of ∼1.5, consistent with the expected effect of increased gene dosage (**Fig. 1A, Data S2**). Nevertheless, there was a wide range of expression of the triplicated genes among individuals with and without T21 (e.g., *IFNAR1*, *DYRK1A*, **Fig. 1B**). Importantly, the transcriptome analysis identified thousands of mRNAs encoded elsewhere in the genome that were also consistently dysregulated in DS (e.g., *MYD88*, *COX5A*, **Fig. 1A-B**, **Data S2**). Gene set enrichment analysis (GSEA) of the transcriptome changes associated with T21 extended previous observations demonstrating activation of the IFN transcriptional response in DS (*6*). In fact, among the top 10 gene sets significantly enriched in T21, seven correspond to IFN signaling and inflammatory pathways (**Fig. 1C, Data S2**). To define which genes on HSA21 associate with specific signaling pathways dysregulated in DS, we correlated their mRNA expression with the rest of the transcriptome using only T21 samples, and then analyzed the resulting matrices of Spearman rho values by GSEA (see **Methods, Fig. S1B**). Importantly, most HSA21 genes had negative correlations with gene signatures of inflammation, with very few mRNAs having consistent significant positive correlations, including the four *IFNRs* and recognized IFN-stimulated genes (ISGs) encoded on HSA21, such as *MX1* and *MX2* (**Fig. S1C-D**, **Data S2**). For example, whereas expression of *IFNAR1* positively correlated with multiple inflammatory pathways (**Fig. 1D, Data S2**), *DYRK1A* expression did not, correlating instead with other pathways (**Fig. 1E, Data S2**). Multiple ISGs not encoded on HSA21 (e.g., *MYD88*, *STAT3, TRIM25)* showed strong positive correlations with *IFNRs* but not with most HSA21 genes (**Fig. 1C-F, Fig. S1C, Data S2**). In contrast, genes in the Oxidative Phosphorylation signature elevated in DS were negatively correlated with *IFNR* expression (e.g., *COX5A*, **Fig. 1C, F, Fig. S1C, Data S2**).

**Fig. 1.**
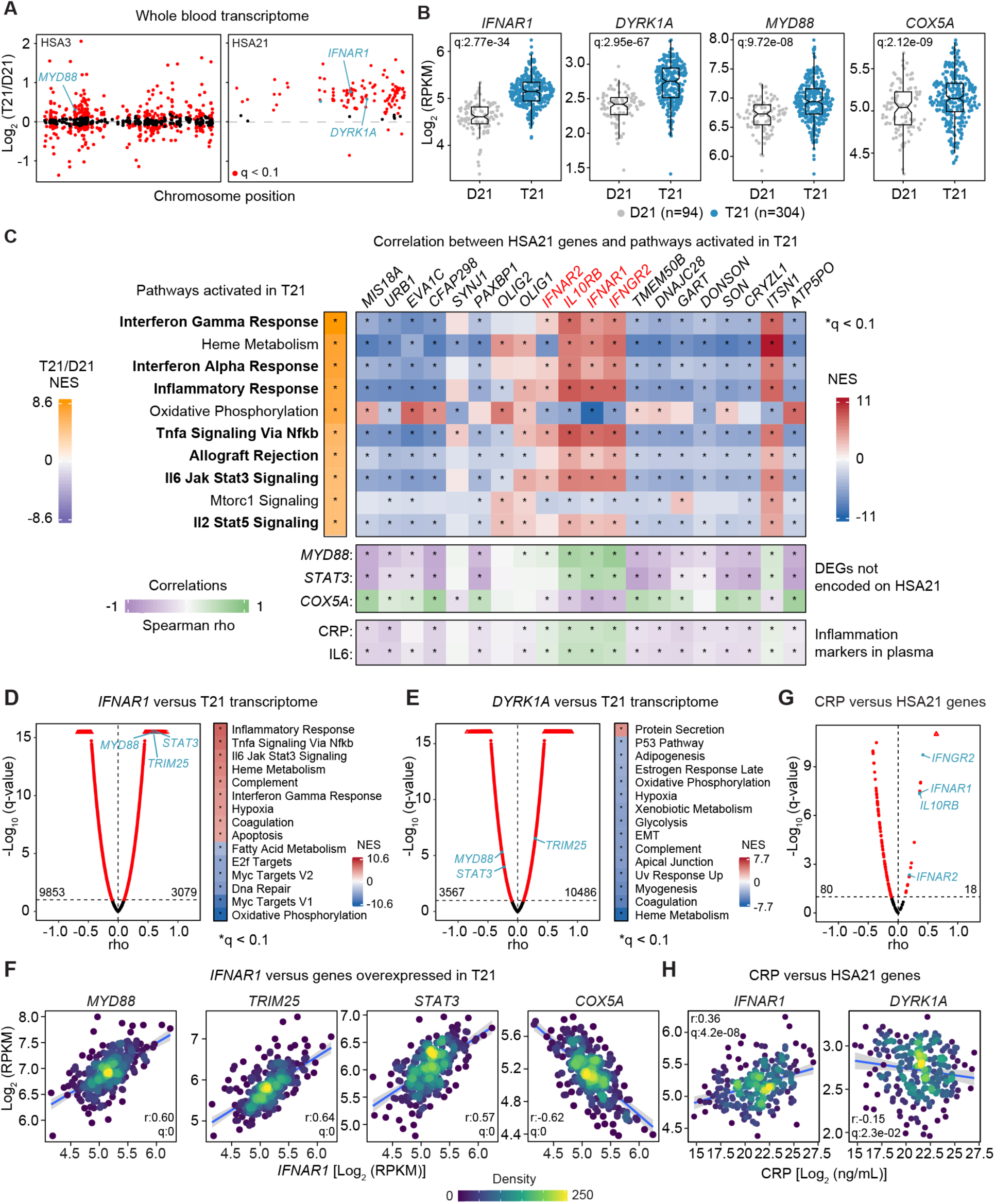
Overexpression of *IFNRs* associates with inflammatory signatures in Down syndrome. **(A)** Manhattan plots of human chromosome 3 (HSA3) and 21 (HSA21) displaying the results of whole blood transcriptome analysis of 304 individuals with trisomy 21 (T21) compared to 96 euploid controls (D21). Differentially expressed genes (DEGs, q<0.1) identified by DESeq2 analysis are labeled in red. **(B)** Sina plots for representative DEGs. **(C) Left**: Heatmap displaying the results of gene set enrichment analysis (GSEA) of transcripts differentially expressed in the whole blood of individuals with T21 compared to D21 controls. Only the top 10 positively enriched pathways are shown. Bold indicates gene signatures indicative of IFN activation and inflammation. NES: normalized enrichment score. **Right top**: Heatmap representing results of GSEA analysis of transcriptome signatures associated with expression of individual HSA21 genes. Spearman correlations were defined between specific HSA21-encoded transcripts and the rest of the transcriptome using only T21 samples. **Middle:** Spearman correlations between transcripts encoded on HSA21 versus the indicated DEGs encoded elsewhere in the genome among individuals with T21. **Bottom:** Spearman correlations between transcripts encoded on HSA21 and the indicated inflammatory markers CRP and IL6 measured in the plasma of individuals with T21. **(D-E)** Volcano plots of Spearman correlations for (D) *IFNAR1* or (E) *DYRK1A* transcript abundance versus the entire transcriptome among individuals with T21. Heatmaps display the results of GSEA of the Spearman correlations. **(F)** Scatter plots showing correlations between *IFNAR1* transcript abundance and DEGs encoded elsewhere in the genome among individuals with T21. **(G)** Volcano plot of Spearman correlations for CRP protein levels in plasma versus transcripts encoded on HSA21 among individuals with T21. **(H)** Scatter plots showing Spearman correlations for CRP versus two transcripts encoded on HSA21. In C-H, statistical significance was determined using GSEA or Spearman correlations after multiple hypothesis correction with Benjamini-Hochberg method, 10% FDR (*q <0.1).

Next, we defined correlations between circulating protein levels of the general inflammatory marker C-reactive protein (CRP) and the pro-inflammatory cytokine IL6 versus expression of individual HSA21 genes among people with T21. Once again, expression of only a small fraction of HSA21 genes positively correlated with CRP and IL6 levels, including the four *IFNRs* (**Fig. 1C, G, Fig. S1E, Data S3**). For example, whereas mRNA expression for *IFNAR1* correlates positively with circulating levels of CRP and IL6, expression of *DYRK1A* is negatively correlated with both immune markers (**Fig. 1H, Fig. S1F**).

Altogether, these results indicate that the heightened inflammatory state observed in individuals with DS is unlikely to be a general effect of the aneuploidy, being instead associated with overexpression of a small subset of specific genes on HSA21, including all four *IFNRs*.

### Normalization of *Ifnr* gene dosage in a mouse model of Down syndrome

The B6.129S7-Dp(16Lipi-Zbtb21)1Yey/J mouse model, herein “Dp16”, carries a segmental duplication of mouse chromosome 16 (MMU16) causing triplication of ∼120 protein-coding genes orthologous to those on HSA21, including the gene cluster of four *Ifnrs* (*15, 16*). Dp16 mice display key phenotypes of DS including increased prevalence of heart defects, craniofacial anomalies, developmental delays, cognitive impairment, a dysregulated antiviral response, and hyperactive IFN signaling (*16–21*). Importantly, gene triplication in the Dp16 model of DS is driven by segmental duplication rather than a freely segregating extra chromosome, akin to DS caused by Robertsonian translocations of HSA21, indicating that Dp16 phenotypes cannot be explained by mere aneuploidy (*16*).

To define if increased gene dosage of *Ifnrs* contributes to DS phenotypes, we used CRISPR/Cas9 genome editing to concurrently knock out all four *Ifnrs*. Given that all four *Ifnrs* employ JAK/STAT signaling and that overexpression of each of them associates with similar inflammatory signatures in our transcriptome analysis, thus creating the potential for genetic redundancy, we designed a strategy to delete the entire 192 kb genomic segment on MMU16 encoding all four *Ifnrs* in wildtype (WT) C57BL/6 mice (see **Methods,** **Fig. 2A, Data S4**). Knockout was confirmed in potential founders by PCR (**Figs. S2A-B**) and Sanger sequencing (**Fig. S2C**). Whole genome sequencing (WGS) confirmed the deletion and did not reveal any other genomic alterations (**Fig. 2A, Fig. S2D**). Heterozygous progeny of this strain (WT^1xIFNRs^) was then intercrossed with Dp16 to normalize *Ifnr* copy number from three to two in a portion of Dp16 offspring (Dp16^2xIFNRs^) (**Fig. S2E**). Expectedly, mRNA expression of all four *Ifnrs* was consistently elevated in Dp16 relative to WT littermates, but this overexpression was corrected in Dp16^2xIFNRs^ mice (**Fig. 2B**). IFNR protein expression was significantly elevated in the surface of bulk white blood cells and multiple myeloid and lymphoid immune cell lineages in Dp16 mice, but not in Dp16^2xIFNRs^ mice (**Fig. 2C-D**, **Fig. S3A-E**). Therefore, Dp16^2xIFNRs^ mice provide an experimental model to define the contribution of *Ifnr* triplication relative to the other ∼120 genes triplicated in the Dp16 animal model of DS.

**Fig. 2.**
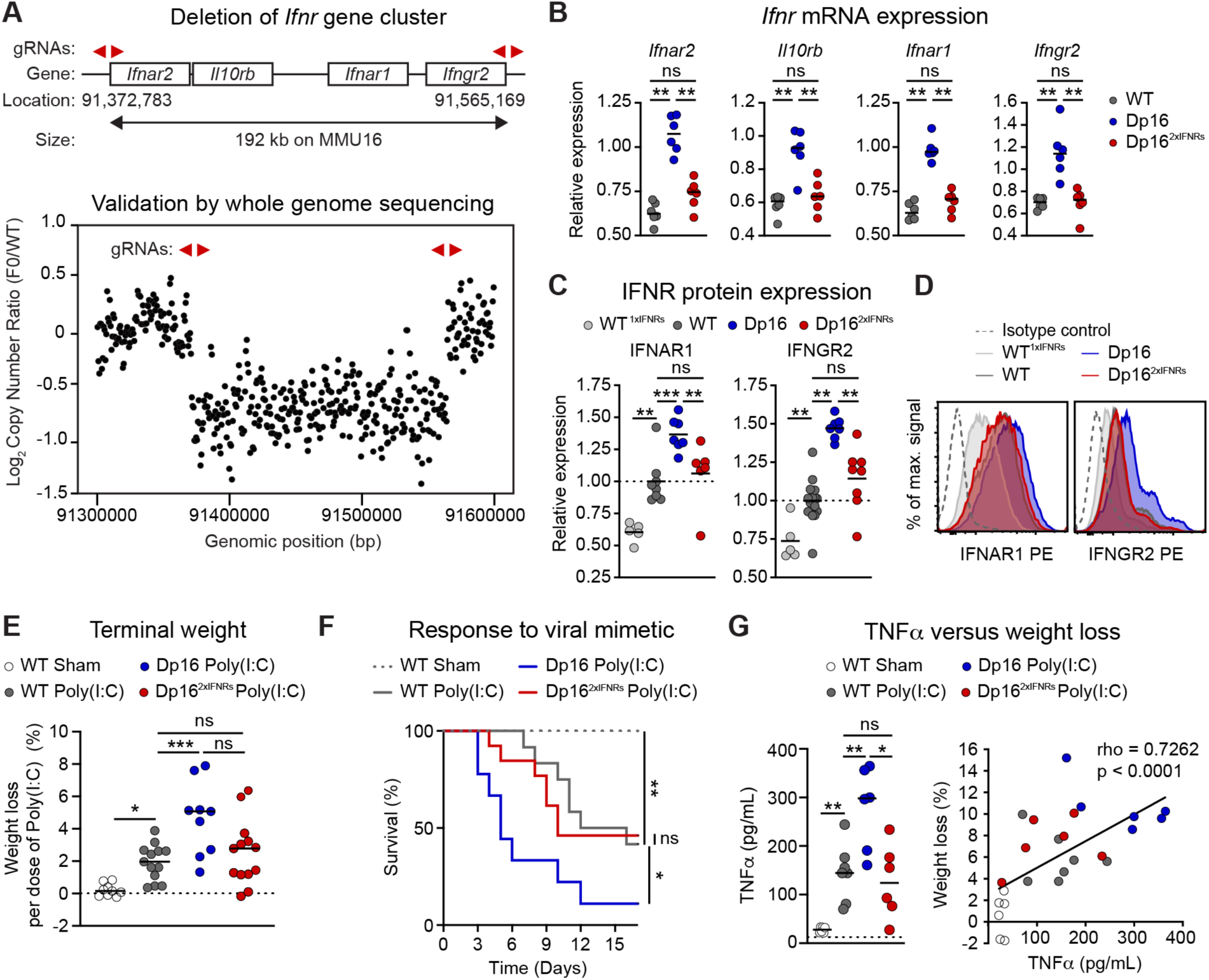
Triplication of *Ifnr* genes drives increased IFNR expression and exacerbated antiviral responses in a mouse model of Down syndrome. **(A) Top:** Diagram indicating genomic locations of the murine *Ifnr* gene cluster on mouse chromosome 16 (MMU16) and guide RNAs (gRNAs, red arrowheads) employed for genome editing using CRIPSR/Cas9 technology. Positions are indicated in base pairs from the GRCm38 annotation of the *M. musculus* genome. **Bottom:** Copy number variant analysis from whole genome sequencing for a candidate founder (F0) bearing a deletion relative to a wildtype (WT) control. **(B)** qRT-PCR analysis of *Ifnr* mRNA expression in hippocampi. Cohorts represent WT controls, the Dp16 mouse model of Down syndrome, and Dp16 normalized for gene dose of just the *Ifnrs* (Dp16^2xIFNRs^) (n=6/group). **(C)** Relative fluorescent intensity (RFI) of geometric mean fluorescent intensities (gMFIs) and **(D)** representative histograms for IFNR protein expression on the surface of CD45+ white blood cells measured by flow cytometry (n=5-16/group). WT^1xIFNRs^ indicates heterozygous *Ifnr* knockouts. **(E)** Results of chronic stimulation with the TLR3 agonist poly(I:C). Mice were injected intra peritoneum with poly(I:C) every other day for 16 days. Percentage of weight lost was normalized to total number of injections (n=9-13/group). **(F)** Kaplan-Meier analysis of survival throughout poly(I:C) treatment (n=9-13; significance by Mantel-Cox log rank test). **(G) Left:** TNFα levels in serum on day 3 of poly(I:C) treatment (n=6-7/group). **Right:** Spearman correlation of TNFα concentration and percent weight lost by each animal on day 3 of poly(I:C) exposure (n=31, line of best fit by simple linear regression). In B, C, E, and G, each dot represents an independent biological replicate with mean indicated and significance determined by Mann-Whitney test. In B-C and E-G, significance is indicated as *p≤0.05, **p≤0.01, ***p≤0.001, and ****p≤0.001.

### Triplication of *Ifnrs* mediates an exacerbated immune response

We previously demonstrated that peripheral immune cells from Dp16 mice are hypersensitive to IFNα and IFNγ stimulation (*21*). Furthermore, upon chronic exposure to the viral mimetic polyinosinic:polycytidylic acid [poly(I:C)], a TLR3 agonist that induces IFNs, Dp16 mice experience exacerbated weight loss and death (*21*). This immune hypersensitivity phenotype can be rescued by pharmacological inhibition of JAK1, a protein kinase that mediates signaling for all three types of IFNs and other cytokines (*21*). We thus tested the impact of reduced *Ifnr* gene dosage on these phenotypes. When stimulated *ex vivo* with IFNα or IFNγ, white blood cells from Dp16 mice show significantly elevated levels of phospho-STAT1 relative to cells from WT mice, but this phenotype is rescued in Dp16^2xIFNRs^ mice (**Fig. S3F-G**). During chronic poly(I:C) challenge, Dp16 mice lost significantly more weight than WT littermates and had to be removed from the experiment much earlier at the human endpoint of 15% weight loss; however, Dp16^2xIFNRs^ mice did not differ from controls in weight loss or overall survival (**Fig. 2E-F**). Analysis of cytokine induction following poly(I:C) treatment revealed significant overproduction of TNFα in Dp16 relative to WT controls, but this was not observed in Dp16^2xIFNRs^ (**Fig. 2G, Fig. S3H**). TNFα is known to mediate inflammation-driven cachexia (*22*), and its levels correlated with weight loss in our experimental paradigm (**Fig. 2G**).

Altogether, these results indicate that triplication of the *Ifnr* cluster results in increased levels of expression of all four *Ifnrs*, leading to dysregulated immune responses.

### Normalization of *Ifnr* gene dosage rescues heart malformations

Congenital heart defects (CHDs) are more common in DS, afflicting around half of newborns with T21 (*3*). To test if *Ifnr* gene dosage contributes to this phenotype in DS, we evaluated the frequency of heart malformations in WT, Dp16, and Dp16^2xIFNRs^ embryos by sectioning the entire developing heart for histological evaluation at embryonic day (E)15.5 (**Fig. 3A-C, Fig. S4A**). In agreement with previous reports (*16, 23, 24*), Dp16 mice displayed significantly elevated frequency of atrial septal defects (ASD) and ventricular septal defects (VSDs), often both types of septal defects at once. Remarkably, this phenotype was corrected in Dp16^2xIFNRs^ mice, which did not present elevated rates of septal defects compared to WT littermates (**Fig. 3D**). To define if JAK/STAT signaling is modulated by *Ifnr* gene dosage in the developing heart tissue, we measured levels of phospho-STAT1 by Western blot. Indeed phospho-STAT1 levels are elevated in the heart tissue of Dp16 mice relative to WT littermates, but this elevation in JAK/STAT signaling is recued in Dp16^2xIFNRs^ embryos (**Fig. S4B**). These results indicate that *Ifnr* triplication disrupts normal heart development, even in the absence of obvious immune stimuli.

**Fig. 3.**
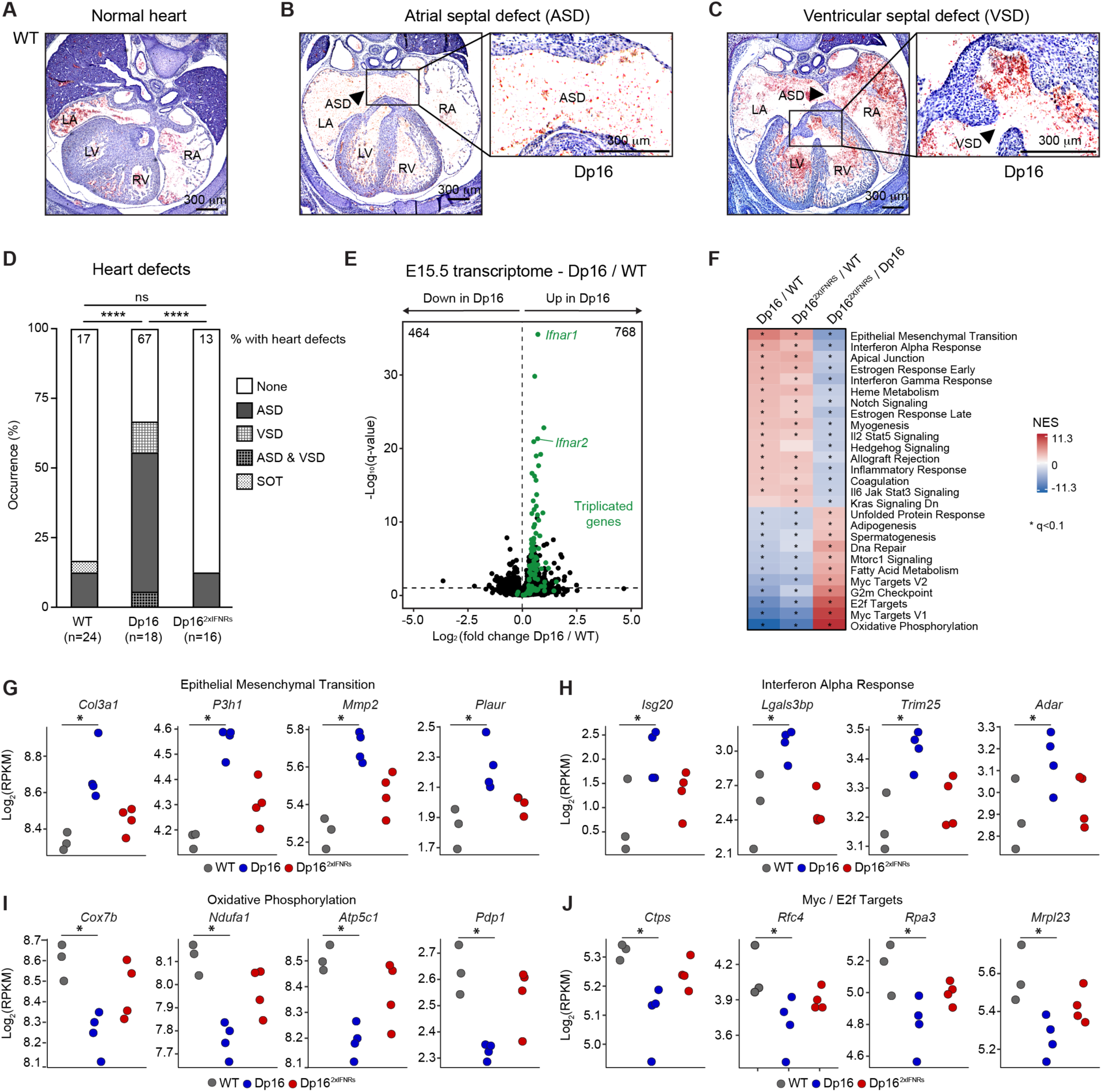
Triplication of *Ifnrs* is necessary for increased incidence of heart malformations in a mouse model of Down syndrome. **(A-C)** Representative images of hematoxylin and eosin staining of serial sections cut through the entire murine heart at embryonic day (E)15.5, including images of normal septation of the four heart chambers in a WT embryo (A), an atrial septal defect (ASD) observed in the Dp16 model of Down syndrome (B), and a ventricular septal defect (VSD) in Dp16 (C). R: right, L: left, A: atrium, and V: ventricle. **(D)** Frequency of heart malformations at E15.5. A single outflow tract (SOT) anomaly is denoted in a WT embryo. Dp16^2xIFNRs^ indicate Dp16 mice normalized for copy number of only the *Ifnr* genes. Significant differences by genotype in total frequency of heart defects was determined by Fisher’s exact test where *p≤0.05 and ****p≤0.0001. **(E)** Volcano plot displaying the results of transcriptome analysis of the developing heart at E15.5, comparing gene expression in the Dp16 transcriptome relative to WT controls, with significant differences determined by DESeq2, 10% FDR (*q<0.1, n=3-4/group). Genes encoded in the triplicated region of MMU16 in Dp16 are labeled green. **(F)** Top gene signatures identified by gene set enrichment analysis (GSEA) of non-triplicated differentially expressed genes in the indicated comparisons. NES: normalized enrichment scores **(G-J)** Dot plots of representative transcripts from major signaling pathways dysregulated in Dp16 and normalized in Dp16^2xIFNRs^ (*q<0.1 by DESeq2).

In order to investigate potential mechanisms underlying this phenomenon, we completed transcriptome analysis of heart tissue at E15.5, which identified ∼1200 differentially expressed genes (DEGs, q<0.1 by DESeq2) in Dp16 relative to WT controls, with only 73 of these DEGs being encoded on the triplicated region (**Fig. 3E; Data S5**). GSEA of DEGs encoded outside of the triplicated region revealed enrichment of genes involved in Epithelial to Mesenchymal Transition (EMT), IFN responses and other immune signatures (e.g. IL2/STAT5 signaling, IL6/JAK/STAT signaling), as well as key developmental pathways, such as Myogenesis, Notch Signaling and Hedgehog Signaling (**Fig. 3F, Data S5**). This analysis also revealed downregulation of genes involved in Oxidative Phosphorylation and multiple gene sets associated with cell growth and proliferation (e.g., MYC targets, E2F targets, mTORC signaling), among others (**Fig. 3F, Data S5**). These same pathways were similarly dysregulated in the hearts of Dp16^2xIFNRs^ mice, albeit to a lesser degree. In fact, comparison of Dp16^2xIFNRs^ versus Dp16 transcriptomes revealed significant normalization of the gene signatures dysregulated in Dp16 (**Fig. 3F**). Examples of EMT genes elevated in Dp16 but normalized in Dp16^2xIFNRs^ include multiple collagen subunits (e.g., *Col3a1*), collagen modifying enzymes (e.g., *P3h1*), matrix metalloproteinases (e.g., *Mmp2*), and the plasminogen activator (*Plaur*) (**Fig. 3G, Data S5**). Canonical ISGs elevated in Dp16 but not in Dp16^2xIFNRs^ include *Isg20*, *Lgals3bp*, *Trim25*, and *Adar* (**Fig. 3H**). Hallmark genes involved in Oxidative Phosphorylation decreased only in Dp16 include subunits of the cytochrome C oxidase complex (e.g., *Cox7b*), the NADH:Ubiquinone Oxidoreductase complex (e.g., *Ndufa1*), the ATP synthase complex (e.g., *Atp5c1*), and the pyruvate dehydrogenase complex (e.g., *Pdp1*) (**Fig. 3I**). Genes involved in cell growth and proliferation depleted only in Dp16 include enzymes involved in nucleic acid synthesis (e.g., *Ctps*), DNA replication factors (e.g., *Rfc4*, *Rpa3*), and mitochondrial ribosomal proteins (e.g., *Mrpl23*) (**Fig. 3J**).

Altogether, these results indicate that triplication of *Ifnrs* elicits a signaling cascade in the developing heart involving elevated JAK/STAT signaling, dysregulation of EMT processes, along with decreased cell growth and proliferation, all of which could contribute to heart malformations.

### Triplication of *Ifnrs* delays development and impairs cognitive function

Children with T21 and Dp16 neonates exhibit delays in achieving developmental milestones (*3, 20*). Relative to WT controls, Dp16 neonates show reduced chance of success in achieving the surface righting reflex as well as ear twitch and auditory startle sensitivities on any given day, but no differences in eye opening (**Fig. 4A**). When females and males are analyzed separately, this phenotype is clearly sexually dimorphic, with Dp16 females showing the most pronounced differences (**Fig. S5A-B**). Notably, Dp16^2xIFNRs^ mice show rescue of all three Dp16 developmental delays (**Fig. 4A**, **Fig. S5A-B**).

**Fig. 4.**
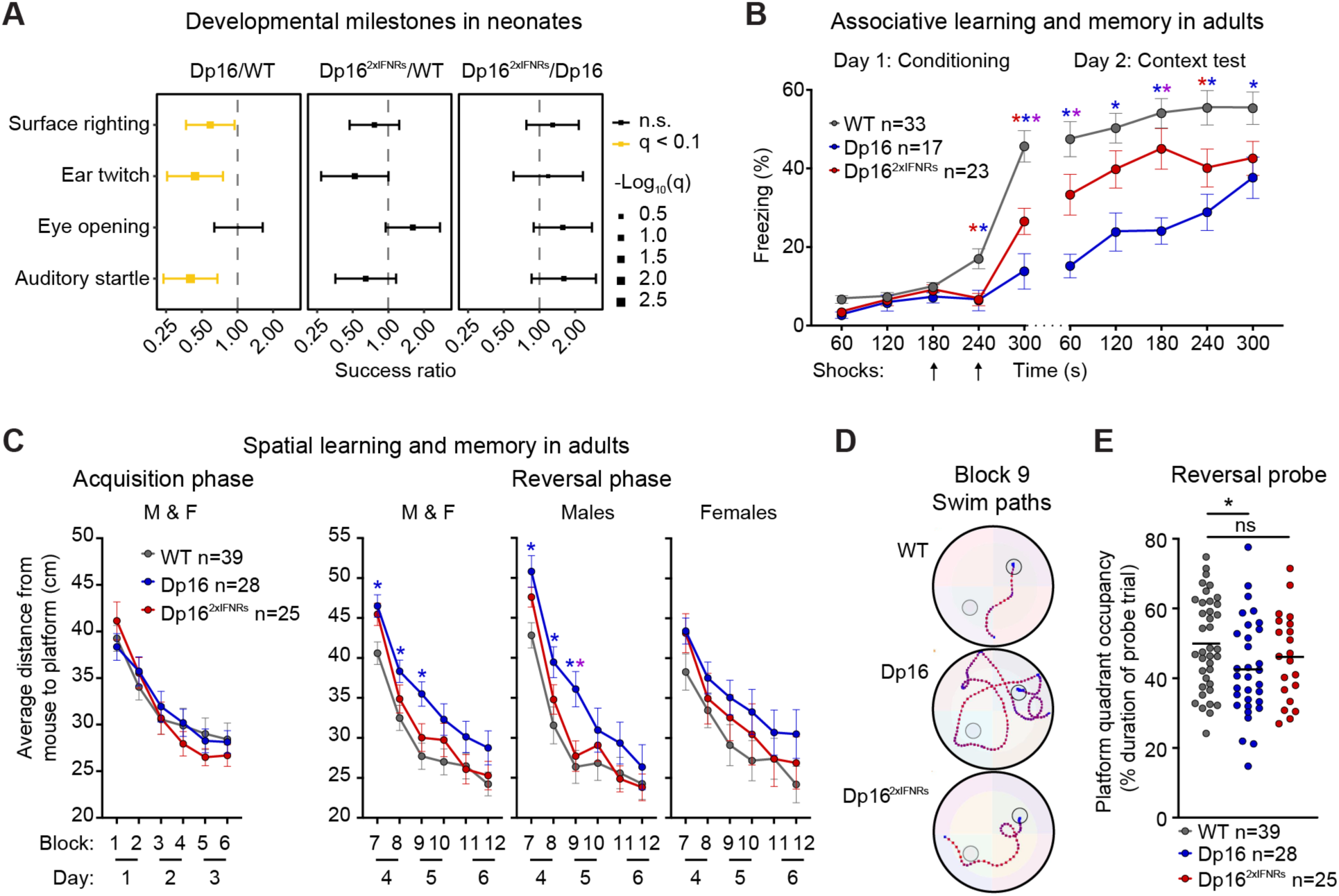
Triplication of *Ifnr*s promotes developmental delays and cognitive deficits in a mouse model of Down syndrome. **(A)** Success ratio of developmental milestone achievement in neonates assessed by mixed effects Cox regression with adjustment for gender and litter and significance set at 10% FDR by Benjamini-Hochberg method. Data are presented as success ratios with label size proportional to q-value and 95% confidence intervals for each pair-wise genotype comparison of the Dp16 mouse model of trisomy 21 (n=32-34, 19-20 male), Dp16 normalized for gene dose of just the *Ifnrs* (Dp16^2xIFNRs^; n=27-30, 16-20 male), and wildtype (WT) controls (n=43-46, 19-20 male). **(B)** Freezing behavior during the contextual fear conditioning test for adult WT (n=33, 13 male), Dp16 (n=17, 8 male), and Dp16^2xIFNRs^ (n=23, 11 male) mice. **(C)** Swim path efficiency of mice navigating to escape platform in the Morris Water Maze (MWM) for male and female (M & F), WT (n=39, 19 male), Dp16 (n=28, 13 male), and Dp16^2xIFNRs^ (n=25, 12 male), during acquisition and reversal phases. **(D)** Representative swim trials for males from block 9 in the MWM reversal phase with platform location denoted for both the acquisition (grey) and reversal (black) phases. **(E)** Mouse occupancy of target quadrant during the reversal probe trial of MWM. In B-D all data represent averages +/- SEM for mice 4-5 months in age and significance by two-way repeated measures (B-C) or one-way ANOVA with post-hoc Tukey’s HSD (E) (*p≤0.05). Significant differences between pairs are denoted by blue * for Dp16 versus WT, red * for Dp16^2xIFNRs^ versus WT, and purple * for Dp16 versus Dp16^2xIFNRs^.

Cognitive deficits in adult mice were then evaluated using contextual fear conditioning (CFC) and Morris water maze (MWM) (*25, 26*). During conditioning in the CFC test for associative learning and memory, mice were presented with two mild foot shocks. Upon the second shock, Dp16 displayed a significantly decreased freezing response relative to WT controls (**Fig. 4B**). When reintroduced to the shock context on day 2, both WT and Dp16 froze at a higher baseline rate relative to the beginning of day 1, yet WT mice froze at significantly higher rate than Dp16. Throughout the experiment, both on day 1 and day 2, Dp16^2xIFNRs^ mice displayed a significant rescue of all phenotypes measured by the CFC test (**Fig. 4B**).

Upon examination of spatial learning and memory via MWM (**Fig. S5C-D**) (*26*), adult mice of all genotypes were equally capable of learning to escape the maze during the acquisition learning phase, as evidenced by decreasing latency and total distance traveled to platform over successive swimming blocks (**Fig. S5E**). However, Dp16 but not Dp16^2xIFNRs^ males swam significantly closer to the periphery when introduced to the maze (**Fig. S5F**). Although this behavior is associated with hindrance of learning (*27*), such thigmotaxis in Dp16 males was moderate, and they still learned to escape the maze as quickly as the other genotypes during acquisition phase (**Fig. S5E**). Immediately upon change of platform location in the reversal phase, both Dp16 and Dp16^2xIFNRs^ presented with deficits in memory extinction (**Fig. S5E**, block 7). However, only Dp16 males exhibited impaired relearning of platform location (**Fig. S5E**, blocks 8-9). All cohorts still improved in performance over time in the reversal phase, as measured by decreased latency and total distance traveled to platform over successive blocks (**Fig. S5E**). These subtle yet significant differences by genotype in the reversal phase are in line with previous publications using MWM to study Dp16 deficits in memory extinction and relearning (*16, 18–20*). Next, to pursue differences by genotype in allocentric memory, we evaluated swim path efficiency. This analysis again showed no difference in acquisition learning by genotype, revealing instead a significant deficit in memory extinction and relearning in Dp16, more pronounced in males, with significant rescue of this phenotype in Dp16^2xIFNRs^ mice (**Fig. 4C-D**). Furthermore, Dp16 but not Dp16^2xIFNRs^ demonstrated significantly reduced target quadrant occupancy during the reversal probe trial (**Fig. 4E**). Lastly, impaired Dp16 motor coordination measured by the rotarod performance test (*28*) was not significantly rescued in Dp16^2xIFNRs^ (**Fig. S5G**).

Altogether, these results indicate that *Ifnr* gene dosage impacts multiple neurological hallmarks of DS, including key early developmental milestones, as well as major domains of cognitive function later in development, including associative learning and memory and spatial memory.

### Normalization of *Ifnr* copy number reduces craniofacial anomalies

Craniofacial abnormalities are a developmental feature of all individuals with DS, with strong inter-individual variation, including brachycephaly, maxillary deficiency, and smaller cranial base (*29*). Given that Dp16 mice reproduce aspects of the distinct craniofacial morphology of individuals with T21 (*17*), we evaluated the impact of *Ifnr* gene dosage on Dp16 skull size and shape. Euclidean Distance Matrix Analysis (EDMA) was performed using 30 craniofacial and mandibular landmarks [LMs, (*30*)] that were collected on 3D-rendered micro-computed tomography (μCT) scans (**Fig. 5A-B**, **Fig. S6A-B**, **Data S6**). 58% (149/259) of all inter-LM distances differed between Dp16 and WT controls, consistent with prior morphometric studies in Dp16 mice (*17*) (**Fig. 5B**, **Fig. S6B**). Remarkably, 23% (34/149) of these differences were normalized in Dp16^2xIFNRs^ mice, including 79% (11/14) of Dp16 mandibular phenotypes (**Fig. 5B**, **Fig. S6B, Data S6**). The remaining inter-LM differences that persisted between controls and Dp16^2xIFNRs^ were less drastic than observed in Dp16 (**Fig. S6C, Data S6**). Key examples include a shortening of the basisphenoid (BS) bone (LMs 24-27), increase in the intersphenofrontal width (a proxy for inter-temple width; LMs 21-22), and alterations in many mandibular inter-LM distances (e.g., LMs 18-20), all of which are observed in Dp16 but attenuated in Dp16^2xIFNRs^ (**Fig. 5C**). These morphometric analyses not only confirm previously reported craniofacial characteristics of Dp16, such as shorter widened skulls (*17*), but also demonstrate substantial effect of *Ifnr* gene dosage on craniofacial morphology.

**Fig. 5.**
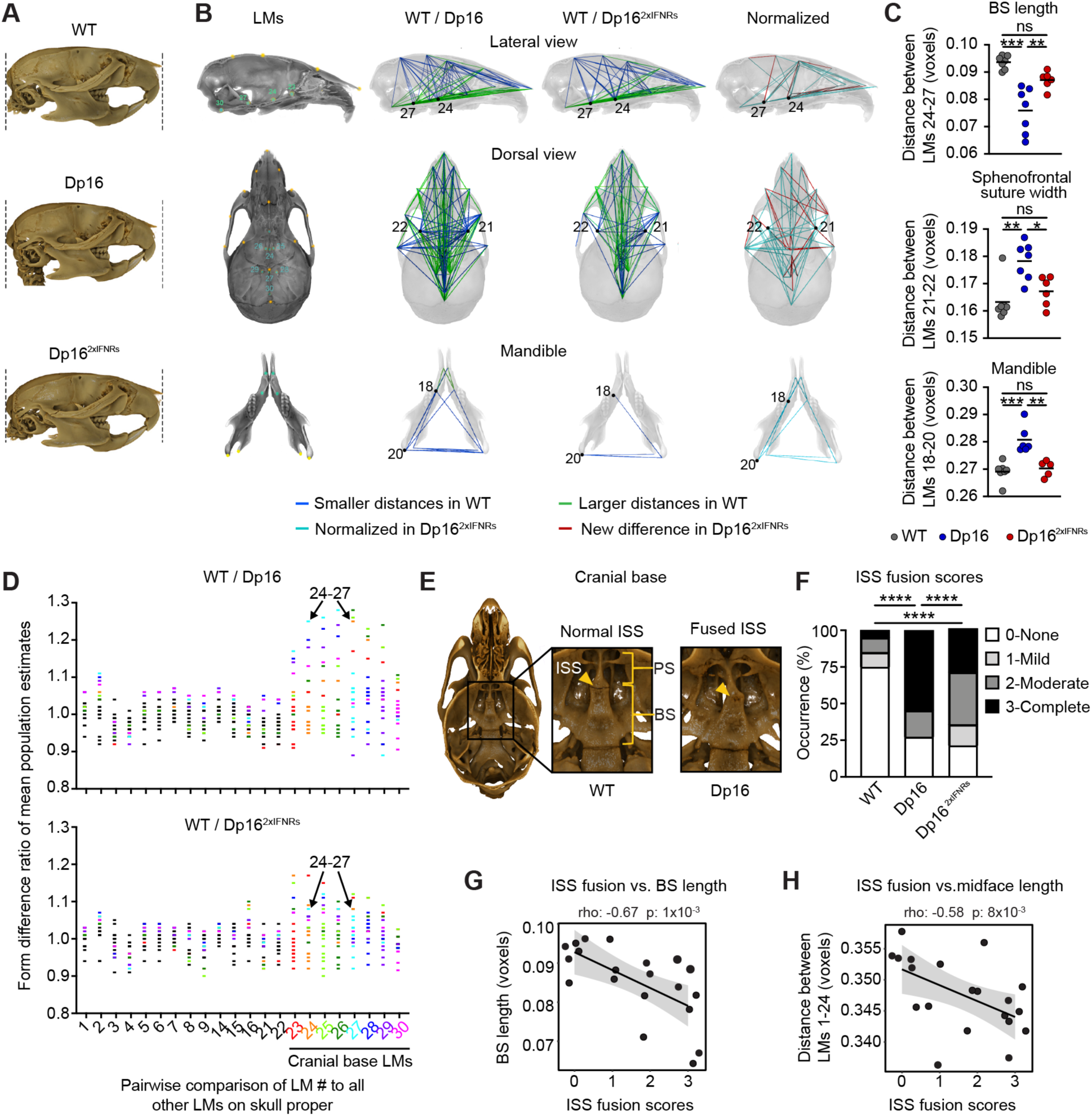
Triplication of *Ifnr*s exacerbates craniofacial anomalies in a mouse model of Down syndrome. **(A)** Representative micro-computed tomography (μCT) images of skulls obtained from wildtype (WT) mice, the Dp16 mouse model of Down syndrome, and Dp16 normalized for gene dosage of just the *Ifnrs* (Dp16^2xIFNRs^). Skulls are aligned and scaled to size based on the same 3D linear measurement. (**B**) Lateral (top) and dorsal (middle) views of the outer portion of the skull proper transparently overlaid over cranial base interior from a WT mouse. Landmarks (LMs) on skull proper (top and middle) and mandible (bottom) are yellow while interior LMs on the cranial base are turquoise. Smaller and larger inter-LM distances (blue and green lines, respectively) in WT relative to Dp16 or Dp16^2xIFNRs^ mice were calculated by Euclidean Distance Matrix Analysis (EDMA, n=6-7 mice/group followed by bootstrapping 10,000x). Inter-LM distances that differed in WT relative to Dp16 that are no longer different in Dp16^2xIFNRs^ (i.e., normalized) are turquoise while inter-LM distances that were not different in WT relative Dp16 that are different in Dp16^2xIFNRs^ are red. **(C)** Scatter plots of inter-LM distances different in Dp16 but not Dp16^2xIFNRs^ relative to WT mice (n=6-7/group; significance by one-way ANOVA with post-hoc Tukey’s HSD). BS: basisphenoid bone. **(D)** Form difference ratio of mean population estimates for inter-LM distances of the skull proper after bootstrapping with color assigned to any inter-LM distance that includes a LM of the cranial base. (**E**) Interior view of cranial base with intersphenoid synchondrosis (ISS), presphenoid (PS) and BS bones labeled on μCT-derived rendered images representative of normal (left) and completely fused (right) ISS. (**F**) Severity of ISS fusion by genotype. Frequency of complete fusion (black) compared between cohorts by Fisher’s exact test (n=11-20). **(G-H)** Spearman correlation of ISS fusion scores against BS (G) or midface length (H) for each skull (n=20/group, with n=6-7 from each genotype). In C and G-H, significant when *p≤0.05, **p≤0.01, ***p≤0.001, and ****p≤0.001.

Many of the differences in inter-LM distances observed in the skulls of Dp16 mice (*31*) involved LMs at the cranial base (i.e., LMs 23 to 30), which in turn were among the most sensitive to *Ifnr* gene dosage (**Fig. 5D, Data S6**). Qualitative inspection of the cranial base revealed a loss of intersphenoidal synchondrosis (ISS) in Dp16 mice, likely due to premature fusion of the presphenoid and BS bones (**Fig. 5E**). This phenotype resembles the early spheno-occipital synchondrosis mineralization observed in individuals with DS (*32*), and its severity was significantly reduced in Dp16^2xIFNRs^ mice (**Fig. 5F**). Early mineralization of the anterior cranial base restricts cranial base and midfacial outgrowth and is often associated with altered calvarium shape (*33*). Notably, Dp16 mice display significantly shortened BS length and midface length (i.e., LMs 24-27 and 1-24, respectively), but these phenotypes are ameliorated in Dp16^2xIFNRs^ (**Fig. S6B**). Furthermore, we observed a significant inverse correlation between basisphenoid length and midface length versus ISS fusion (**Fig. 5G-H**).

Altogether, these results indicate that triplication of *Ifnrs* contributes to major craniofacial features distinctive of DS, further supporting a role for hyperactive IFN signaling in the dysregulated development of diverse organ systems in DS, including skeletal morphogenesis.

## Discussion

Despite significant research efforts, the mechanisms by which T21 causes the developmental and clinical hallmarks of DS remain poorly understood (*3*). In addition to the possibility that multiple genes contribute to a specific phenotype, the aneuploidy itself could exert effects independent of gene content (*12*). Clearly, elucidation of gene-phenotype relationships in DS would accelerate therapeutic strategies to ameliorate the ill effects of the trisomy. Within this framework, deciphering the mechanisms by which T21 causes lifelong IFN hyperactivity and dysregulation of downstream signaling pathways could enable immunomodulatory strategies to improve health outcomes in DS.

Using whole blood transcriptome analysis in a large cohort of individuals with DS, we observed that overexpression of only a small subset of genes encoded on HSA21 correlates with gene signatures indicative of IFN hyperactivity and inflammation, including the four *IFNRs*. This exercise demonstrated that overexpression of individual triplicated genes can be tied to dysregulation of specific pathways in DS. Although hyperactive IFN signaling has been noted in cells with T21 since the 1970’s, the contribution to key phenotypes of DS has not been defined (*34*). In the Ts16 mouse strain carrying triplication of essentially all MMU16 genes through a centric-fusion translocation, including many genes not orthologous to HSA21, reduction of IFN signaling improved some aspects of Ts16 fetal development (*34*). However, because these mice die shortly after birth, examination of postnatal phenotypes was not feasible (*34*). We therefore employed genome editing to test the impact of normalizing *Ifnr* dosage in the Dp16 preclinical mouse model of DS bearing a segmental duplication involving ∼120 protein coding genes on MMU16 orthologous to HSA21 (*16, 35*). This approach revealed that *Ifnr* triplication contributes to a dysregulated antiviral response, septal heart malformations, developmental delays, cognitive deficits, and craniofacial abnormalities in mice. These results expand and strengthen an increasing body of work documenting harmful effects of aberrant IFN signaling in human development (*13*), while supporting the notion that DS can be understood in part as an interferonopathy (*6, 14, 36*).

Our results demonstrating that *Ifnr* triplication underlies an exacerbated immune response may help explain the high rate of morbidity and mortality from respiratory infections observed in DS, as well as the increased rate of autoimmune disorders (*4, 5, 37*). T21 is a top risk factor for severe COVID-19, leading to greatly increased rates of hospitalization and mortality (*38, 39*). IFN signaling plays multiple roles in COVID-19 pathology, with both protective and harmful effects being documented. The protective effects of IFN signaling are demonstrated by studies showing decreased IFN signaling associated with severe symptomatology (*40–42*), the presence of autoantibodies blocking IFN signaling (*43–47*), and genetic variants that impair IFN signaling (*48, 49*). However, reduced *Ifnar1* copy number prevents lung pathology in mouse models of both SARS-CoV-1 and SARS-CoV-2 infections (*50, 51*). Furthermore, both Type I and Type III IFNs contribute to disruption of lung barrier function during viral infections (*52, 53*). Regarding autoimmunity, IFN hyperactivity has been consistently associated in the general population with development of autoimmune disorders more common in DS, as both pharmacological IFN treatment and genetic variants leading to heightened IFN signaling increase the risk of developing autoimmune conditions (*54, 55*).

Our findings define a role for the *Ifnrs* during embryonic heart development, even in the absence of obvious immune triggers. In mice, numerous large regions orthologous to HSA21 were identified that can contribute to increased rate of heart malformations, some of which include the *Ifnr* gene cluster (**Fig. S7**) (*3, 16, 23, 24, 56, 57*). Notably, single nucleotide polymorphisms in *IFNGR2* and *IL10RB* have been associated with risk of CHD in DS (*58*). Nevertheless, our results are the first demonstration that normalization of *Ifnr* copy number is sufficient to rescue this phenotype. Furthermore, we show that *Ifnr* triplication drives increased JAK/STAT signaling and dysregulation of major signaling pathways in the developing heart, including a gene expression program indicative of reduced cell growth and proliferation. These findings have implications to the understanding of CHD and heart malformations in other contexts. For example, in the typical population, hyperactive IFN signaling has been indirectly implicated in abnormal prenatal heart development during maternal lupus and viral infections (*59, 60*). Although there has been much work to characterize the craniofacial morphology in people with DS and mouse models of T21, our results showing that *Ifnr* gene dosage affects development of the skull proper, cranial base, and mandible provides much needed mechanistic insight about the etiology of this phenotype. Furthermore, these findings have widespread implications for understanding of fetal development beyond DS. For example, maternal virus infections are known to alter the inflammatory milieu for a developing fetus, can disrupt bone development, and cause cranial calcifications and microcephaly, which are characteristics of individuals affected by monogenic interferonopathies (*61*). Lastly, the impact of *Ifnr* triplication on early development and cognitive function lends support to the notion of hyperactive IFN signaling as a driver of brain pathology and abnormal neurodevelopment both after congenital infections and in Type I Interferonopathies (*13, 60*).

These observations justify the testing of immune-modulatory agents in DS. IFN signaling can be attenuated with agents approved for treatment of diverse autoinflammatory conditions, most prominently JAK inhibitors (*21, 62*). JAK inhibition blocks the immune hypersensitivity phenotype observed in Dp16 mice (*21*), and a clinical trial for JAK inhibition in DS is already underway (NCT04246372). In young Dp16 mice, acetaminophen treatment decreased microglia activation and improved cognitive performance (*63*). In alternative mouse models of DS also harboring triplication of *Ifnrs* and displaying hyperactive IFN signaling in multiple brain regions (*20*), developmental delays were rescued by prenatal treatment with the anti-inflammatory natural compound apigenin (*64*), and cognitive performance was improved by treatment with the anti-inflammatory agent minocycline (*65*). An increasing appreciation about the role of maternal and fetal antiviral immunity in development of congenital disease supports the need for additional research to dissect which of the *IFNRs*, their specific ligands, downstream kinases, and subsequent inflammatory pathways are main contributors to the diverse phenotypes of DS. Lastly, how *IFNR* triplication interacts with genetic variants and environmental factors, especially diverse immune stimuli, to potentiate DS phenotypes is yet to be elucidated. In sum, our results demonstrate a role for cytokine signaling in development of both pre- and post-natal phenotypes in DS, supporting the pursuit of anti-inflammatory agents across the lifespan in this vulnerable population.

## Acknowledgments

We thank LF Bush for administrative support, MM Ahmed for advice on brain harvests, as well as H Potter for use of his perfusion machine. We also thank J Gross of the Genetics Core Facility at National Jewish Health (NJH), and S Beard of the University of Colorado (CU) Diabetes Research Center Cell and Tissue Analysis Core (NIDDK grant P30-DK116073) for technical support in novel mouse model generation and flow cytometry, respectively. We thank T Bruno at the University of Pittsburgh and Ivan D’Orso at the University of Texas Southwestern for laboratory space to process remote HTP samples, J Shaw for consultation on statistics, and DE Clouthier for introduction to TC Cox. We thank vivarium personnel for support with mouse colony maintenance as well as all mice comprising this study. Lastly, we are truly grateful for all participants of the HTP.

## Author contributions

Conceptualization, JME, KDS, KAW, KS, TCC, and RM; Methodology, JLM, KAW, KDT, MDG, KPS, KDS, KS, RM, NB, TCC, and ML; Software, MDG, NPE, and KTK.; Validation: JLM, KAW, KDS, CC, KTK, DJO, and LND; Formal analysis, KAW, NPE, MDG, KTK, CC, KDS, KTK, LND, JB, MDG, TCC, and RM; Investigation, JG, KAW, RM, DJO, ZA, RM, JB, DT, LND, KS, CC, KAS, NB, RDA, LLC, HRL, ECB, and PA; Resources, KAW, RM, JLM, JB, DT, HD, KPS, REG, and KAS.; Data curation, NPE, MDG, KTK, KDS, KAW, RDA, LLC, TCC, LLC and RDA; Writing – original draft, KAW, KDS, JME, and TCC; Writing – review & editing: all authors.

## Data and materials availability

All data to evaluate conclusions in this manuscript are present. WGS and RNAseq data were deposited in public databases. The new *Ifnr* knockout mouse strain will be shared upon publication.

## Supplementary Materials for

### Other Supplementary Materials for this manuscript include the following

Data S1 to S6: Data underlying Figs. 1-5 and Figs. S1 to S6.

### Clinical study design

The main goals of the clinical study were to determine baseline changes in circulating inflammatory markers of people with Down syndrome (DS, trisomy 21; T21) in comparison to typical (D21) controls and among individuals with T21. Toward this end, we utilized RNA sequencing (RNA-seq) and multiplexed immunoassays with Meso Scale Discovery (MSD) technology. All human participants (**Data S1**) were enrolled under the Crnic Institute Human Trisome Project protocol approved by the Colorado Multiple Institutional Review Board (COMIRB, protocol number #15-2170, NCT02864108). All procedures were performed in accordance with COMIRB guidelines and regulations. Written informed consent was obtained from participants older than 7 years old who were cognitively able or by guardians of each participant. The HTP cohort analyzed in this study consists of 400 individuals (304 with DS, **Fig. S1A**). None of the participants with DS in the HTP cohort reported signs of active infection for at least 2 weeks prior to blood draw. The HTP enrolls individuals with DS as well as healthy, non-affected family members and unrelated controls. This study was observational and involved collecting participant demographic and health information and various biological samples for biobanking. The banked materials are de-identified and made publicly available to DS investigators for research purposes (trisome.org). The study was conducted in accordance with the Declaration of Helsinki.

### Whole-blood RNA sequencing

RNA-seq was done as previously described (*66*). Briefly, peripheral blood was collected from human study participants in PAXgene RNA Tubes (PreAnalytiX/Qiagen, Cat# 762165). RNA was purified using the PAXgene Blood RNA Kit (Qiagen, Cat# 762164). RNA quality was assessed using an Agilent 2200 TapeStation and quantified on a Qubit fluorometer. Globin RNA depletion, poly-A(+) RNA enrichment, and strand-specific library preparation were carried out using a GlobinClear kit, NEBNext® Poly(A) mRNA Magnetic Isolation Module, and NEBNext Ultra™ II Directional RNA Library Prep Kit for Illumina. Paired end 150 bp sequencing was carried out on an Illumina NovaSeq 6000 instrument by Novogene Co., Ltd. Reads were demultiplexed and converted to fastq format using bcl2fastq (bcl2fastq v2.20.0.422).

RNA-seq data yield was ∼33-103 × 10^6^ raw reads and ∼21-69 × 10^6^ final mapped reads per sample. Data quality was assessed using FASTQC (v0.11.5; RRID:SCR_014583) and FastQ Screen (v0.11.0,; RRID:SCR_000141). Trimming and filtering of low-quality reads was performed using bbduk from BBTools (v37.99; RRID:SCR_016968) (*67*) and fastq-mcf from ea-utils (v1.05; RRID:SCR_005553). Alignment to the human reference genome (GRCh38) was carried out using HISAT2 (v2.1.0; RRID:SCR_0155303) (*68*) in paired, spliced-alignment mode with a GRCh38 index with a Gencode v33 annotation GTF (RRID:SCR_014966), and alignments were sorted and filtered for mapping quality (MAPQ > 10) using Samtools (v1.5) (*69*). Gene-level count data were quantified using HTSeq-count (v0.6.1; RRID:SCR_005514) (*70*) with the following options (--stranded=reverse –minaqual=10 –type=exon -- mode=intersection-nonempty) using a Gencode v33 GTF annotation file. Differential gene expression in T21 versus D21 was evaluated using DESeq2 (version 1.28.1; RRID:SCR_015687)8 in R (version 4.0.1), using q < 0.1 (FDR < 10%) as the threshold for differentially expressed genes. The Whole Blood RNA-seq data have been deposited in NCBI Gene Expression Omnibus.

### MSD cytokine analysis

Peripheral blood was collected into BD Vacutainer® K2 EDTA tubes (BD, 366643), then processed within 2 h of blood draw by centrifugation at 700 x g for 15 min to separate plasma, buffy coat, and red blood cells (RBCs). Two technical replicates were measured for each EDTA plasma sample using the Meso Scale Discovery (MSD) multiplex immunoassay for CRP and IL6 (Cat# K15248D). Assays were performed following manufacturer’s instructions and concentration values were calculated against a standard curve using provided calibrators. Plasma concentration values (pg/mL) for each of the cytokines and related immune factors measured across multiple MSD assay plates was imported to R, combined, and analytes with >10% of values outside of detection or fit curve range flagged. For each analyte, missing values were replaced with either the minimum (if below fit curve range) or maximum (if above fit curve range) calculated concentration and means of duplicate wells used in all further analysis. Cytokine analysis was done of 249 study participants with T21 for which a matched whole blood transcriptome analysis was available.

### Gene set enrichment analysis of clinical samples

GSEA (*71*) was carried out in R using the fgsea package (v 1.14.0; RRID:SCR_020938) (*72*) in R, using Hallmark gene sets (*73*) and either log_2_-transformed fold-changes (for RNA-seq) or Spearman *rho* values (for correlations) as the ranking metric.

### Spearman Correlation Analysis

As previously described (*74*), Spearman *rho* values and p values were calculated among the indicated datasets using the *rcorr* function from the Hmisc package (v 4.4-0), with Benjamini-Hochberg correction of p values and an estimated false discovery fate threshold of 0.1. For visualization, XY scatterplots with points colored by local density were generated using a custom density function and the ggplot2 (v3.3.1; RRID:SCR_014601) package (*75*).

### Generation of mice with deletion of interferon receptor genes on mouse chromosome 16

The 192 kb genomic segment containing just the four interferon receptor (*Ifnr*) genes syntenic to human chromosome 21 (HSA21) was deleted in mice using CRIPSR/Cas9 gene editing as previously described for smaller regions of the genome (*35*). Briefly, CRISPR/Cas9 target sites were identified using http://crispr.mit.edu/ with scores of 84-96 to predict specific deletions. Two guide RNAs (gRNAs) were synthesized per target site flanking the *Ifnr* gene cluster on mouse chromosome 16 (MMU16) **(Data S4, Tab A)**. C57BL/6NTac zygotes (Taconic) were microinjected with Cas9 mRNA and four total gRNAs, then implanted into pseudopregnant females. Mutant mice were made in collaboration with Dr. Jennifer Matsuda and James Gross of the Genetics Core Facility at National Jewish Health, CO.

### Genotyping of mice

DNA from mice was genotyped by polymerase chain reaction (PCR) based on previous protocols (*35*). Briefly, genomic DNA was prepared from 1-2 mm of toe, tail or ear tissue using the HotSHOT method (*76*) then run through PCR according to **Data S4, Tabs B-C** or outsourced for automated genotyping by RT-PCR with specific probes designed for each gene (Transnetyx, Cordova, TN).

### Sanger sequencing of mutant mice

Potential founders (F0s) were genotyped by PCR to identify those that appeared to lack the entire *Ifnr* gene cluster without additional large chromosomal rearrangements such as inversions or duplications on MMU16. These F0 mice were bred to wildtype (WT) C57BL/6N (Taconic) mice to generate heterozygous F1 progeny (herein WT^1xIFNRs^). PCR products spanning the deleted region were generated and subjected to Sanger sequencing using a 3730xl DNA Analyzer (ThermoFisher Scientific) (3) to identify transmission of a single modified allele to progeny. Sequence-verified F1 mice with identical deletion events were then selected to maintain the line.

### Whole genome sequencing and copy number variant analysis of mutant mouse

Whole genome sequencing (WGS) was used to confirm clean chromosomal rearrangements of a proven F0 WT^1xIFNRs^ by copy number variant (CNV) analysis as previously described (*77*). Briefly, peripheral blood was collected in tubes with heparin sodium salt (Sigma, 2106) from the submandibular vein of the F0 male WT^1xIFNR^ and two male C57BL/6NTac WT controls. Red blood cells were then lysed by ammonium chloride-potassium (ACK, (*78*)) then DNA isolated using the AllPrep DNA/RNA/Protein Mini Kit (Qiagen, Cat# 80004).

DNA was quantified using the Qubit 3 Fluorometer according to manufacturer’s instructions (Invitrogen). Libraries were generated from 1 μg DNA using NEBNext® DNA Library Prep Kit and indices were added to each sample. The genomic DNA was randomly fragmented to a size of 350bp by shearing, then DNA fragments were end polished, A-tailed, and ligated with the NEBNext adapter for Illumina sequencing, and further PCR enriched by P5 and indexed P7 oligos. The PCR products were purified (AMPure XP system) and resultant libraries were sequenced on an Illumina NovaSeq 6000 instrument (Novogene, Singapore, China).

Analysis of library complexity and high per-base sequence quality across all reads was performed using FastQC software. Other bioinformatic steps included trimming of low-quality bases, short reads, and adaptor sequences, with the fastqc-mcf tool; read alignment to the UCSC GRCm38 reference genome using BWA; filtering of high-quality mapped reads with SAMtools; and final quality performed using RSeQC. Using the package CNV-seq (RRID:SCR_013357) (4), we confirmed the presence of *Ifnr* deletion by searching for copy number variations (CNVs) between WT and the F0 male WT^1xIFNR^. As we were not interested in discovering novel CNVs, rather confirming that a) our knockout was effective and b) that there were no major off-target effects, we iteratively increased the --minimum-windows-required parameter until no CNVs were called between the WT mice of identical genetic background, leading to a parameter value of 14. Given a significance level of (p), 0.01 and a CNV detection threshold ratio (r) of 0.06, the theoretical minimum window size was determined as using the default method originally described (4). The window size used for the detection of CNVs was 1.5x (default) the theoretical minimum window size. Using these parameters, we confirmed the presence of our intended deletion with no other coherent CNVs of similar size. WGS data were deposited in NCBI SRA.

### Animal Husbandry

All animal experiments were approved by the Institutional Animal Care and Use Committee (IACUC) at the University of Colorado Anschutz Medical Campus, under Protocol #00111. Candidate F1 progeny of the validated F0 WT^1xIFNRs^ were backcrossed to WT C57BL/6J (The Jackson Laboratory) for at least 3 generations before female WT^1xIFNRs^ were intercrossed with males from the Dp16 mouse model of Down syndrome (DS), also of the C57BL/6J background. This Dp(16Lipi-Zbtb21)1Yey/J (hereafter “Dp16”) mouse model of DS (*79*) was originally purchased from The Jackson Laboratory (Cat# JAX:013530, RRID:IMSR_JAX013530) as well as gifted from Drs. Diana Bianchi and Faycal Guedj (National Institutes of Health, NIH) and intermixed and maintained on the C57BL/6J background. After intercrossing female WT^1xIFNRs^ with male Dp16, mice were confirmed to be at least 87.5% C57BL/6J via Transnetyx automated PCR services using single nucleotide polymorphisms from 48 alleles (Cordova, TN) before use in experiments with the remaining mixed background on C57BL/6N (Taconic).

Mice were housed separately by gender in groups of 1-5 mice/cage under a 14:10 light:dark cycle with controlled temperature and had ad libitum access to food (6% fat diet) and water. All animal protocols were reviewed and approved by the Institutional Animal Care and Use Committee at the University of Colorado Anschutz Medical Campus and performed in accordance with the NIH guidelines for the care and use of animals in research.

### qRT-PCR to assess *Ifnr* transcript expression in brain hippocampi

Mice 31-34 weeks of age were euthanized by CO_2_ asphyxiation and cervical dislocation then immediately perfused with 1x PBS using a Perfusion Two Automated Perfusion Instrument (Leica, Cat#39471005). Brains were removed from skulls then cut in half to divide the right and left side of the brain performing a sagittal section. The hippocampus was then manually dissected out of the right brain hemisphere and homogenized in Lysing Matrix D tubes (MP Biomedicals, Cat#6913500) containing 594 μL of lysis buffer RLT Plus (Qiagen, Cat#1048849) and 6 μL of 2-mercaptoethanol (Sigma, Cat#M3148) for 30 s using a Mini-Beadbeater-24 (BioSpec Products, Cat#112011) then frozen at -80°C. Upon quick freeze/thaw in a water bath at 37°C, total RNA was isolated using the AllPrep DNA/RNA/Protein Mini Kit (Qiagen, Cat#80004)). RNA was quantified using the Qubit 3 Fluorometer (Invitrogen) and cDNA generated from 100 ng of RNA using the Applied Biosystems High-Capacity cDNA Reverse Transcription Kit (Cat#43-688-14). qRT-PCR was then performed as previously described (*21*) using the Applied Biosystems Viia7 384-well block real time PCR system. Briefly, qRT-PCR master mix was prepared with Applied Biosystems SYBR Select Master Mix for CFX. Standard curves were run for every primer pair in each qRT-PCR experiment to ensure efficient amplification of target transcripts within all experimental tissues. All samples were run in triplicate, averaged, and normalized to 18s rRNA. Primer sequences are provided in **Data S4, Tab D**.

### Spectral flow cytometry to assess IFNR surface expression on white blood cells

Peripheral blood was collected from the submandibular vein of mice 17-25 weeks of age into tubes of lithium heparin (Sarstedt, Cat#41.1393.105) then stained as previously described with minor alterations (*21*). Briefly, 25 μL of fresh whole blood was pre-incubated with anti-mouse CD16/32 (BioLegend TruSTain FcX, clone 93, Cat#101320) at 1:100 for 10 mins at RT then spiked with a concentrated stain of all surface markers for 30 mins at 4°C.

Staining with fluorochrome-conjugated antibodies purchased from BioLegend, unless otherwise noted, included the following diluted in BD Horizon Brilliant Stain Buffer Plus (BD Biosciences, Cat#566385): mouse CD4-BV711 (clone RM4-5, Cat#100550; RRID:AB_2562607), IA/IE-BV650 (clone M5/114.15.2, Cat#107641; RRID:AB_2565975), Ly6C-BV605 (clone HK1.4, Cat#128036; RRID:AB_2562353), CD8-BV510 (clone 53-6.7, Cat#100752; RRID:AB_2561389), CD115-BV421 (clone AFS98, Cat#135513; RRID:AB_2562667), CD45 (clone 104, Cat#109820; RRID:AB_492872), SiglecF-BB515 (clone E50-2440; BD Horizon, Cat#564514; RRID:AB_2739601), CD11b-AF532 (clone M1/70; Invitrogen, Cat#58-0122-82; RRID:AB_2811905), NK1.1-BB700 (clone PK136; BD Horizon, Cat#566502; RRID:AB_2744491), CD3-PE/Cy7 (clone 17A2, Cat#100220; RRID:AB_312684), Ly6G-BV605 (clone 1A8, Cat#127624; RRID:AB_10645331), B220-AF700 (clone RA3-6B2; Invitrogen, Cat#56-0452-82; RRID:AB_897458), IFNAR1-PE (clone MAR1-5A3, Cat#127312; RRID:AB_284000) or isotype mouse IgG1k-PE (clone MOPC-21, Cat#981804; RRID:AB_2847529), IFNGR2-PE (clone MOB-47, Cat#113603; RRID:AB_313560) or isotype Armenian hamster IgG-PE (clone HTK888, Cat#400907; RRID:AB_326593), IL10RB-PE (REAffinity clone REA856; Miltenyi, Cat#130-114-497; RRID:AB_2726752) or isotype REA control human IgG1-PE (REAffinity clone REA293; Miltenyi, Cat#130-113-428; RRID:AB_2733893).

After staining, whole blood underwent two immediate 2 and 5 min incubations in 200 μL of ACK (*78*). Cells were then washed twice in flow cytometry wash buffer (1x PBS, 2% FBS, 10 mM HEPES pH 7.5, and 0.1% sodium azide) then fixed for 10 mins at RT in paraformaldehyde (PFA; Ted Pella Inc, Cat#18505) diluted to 4% in 1x PBS. Fixative was washed-out in flow cytometry wash buffer then cells analyzed using a five laser Cytek® Aurora spectral flow-cytometer.

### Spectral flow cytometry to assess phopho-STAT1 in white blood cells

Peripheral blood was collected from the submandibular vein of mice 17-25 weeks of age into tubes of lithium heparin (Sarstedt, Cat#41.1393.105) then stained as previously described with the minor alterations described above for IFNR stains (*21*). Briefly, 25 μL of blood were subjected to RBC lysis then stimulated for 30 minutes at 37C with 10,000 units/mL of recombinant IFN-α2A (R&D Systems, Cat# 12100-1) or 100 units/mL of recombinant IFNγ (R&D Systems, Cat# 485-MI). Antibodies conjugated to methanol-stable fluorophores targeting epitopes that are not stable through fixation were also included in the stimulation media in the presence of FcR block (i.e., SiglecF, Ly6C, CD115, CD8, and CD11b). Following stimulation, cells were washed in FACS buffer, subjected to BD Lyse-Fix buffer (BD Biosciences, Cat# 5580249), then to permeabilization buffer III (BD Biosciences, Cat# 558050). Cells were then stained with fluorophore-conjugated antibodies specific for the following epitopes that were not destroyed with fixation: CD45, CD3, CD4, B220, NK1.1, Ly6G, IA/IE, and phospho-STAT1 (Tyr701) (BD Biosciences, Cat# 562069; RRID:AB_399855). Fixative was washed-out in FACS buffer then cells analyzed using a five laser Cytek® Aurora spectral flow cytometer. Additional antibody and buffer information is provided above in the IFNR staining section. Flow cytometry data was similarly analyzed with FlowJo Software (Becton, Dickinson & Company).

### Enzyme-linked immunoassay to assess IFNAR2 protein in serum

Peripheral blood was collected from the submandibular vein of mice 11-36 weeks of age into tubes containing serum gel with clotting activator (Sarstedt, Cat#41.1500.005). Serum was isolated by spinning at 10,000 xg for 5 mins at RT, separated out from the original tube and serum gel, then frozen at -80°C. Upon first freeze/thaw on ice, serum was brought to RT then diluted 1:1000 and analyzed by the sandwich enzyme-linked immunoassay (ELISA) Mouse IFN-alpha/beta R2 ELISA Kit according to manufacturers’ instructions (RayBiotech, Cat# ELH-IFNabR2-1). Notably, both the capture and detection polyclonal antibodies were generated against the full-length murine IFNAR2 protein (RayBiotech Technical Support, personal communication). Plates were analyzed on the Synergy H4 Hybrid Multi-Mode Microplate Reader (BioTek).

### Chronic exposure of mice to a viral mimetic

Experiments with polyinosinic:polycytidylic acid [poly(I:C)] in mice 18-31 weeks old were done as previously described (*21*). Briefly, 10 mg/kg of poly(I:C) (HMW) VacciGrade (InvivoGen, Cat#31852-29-6) was administered intraperitoneally at 2-day intervals for up to 16 days. Animals were sacrificed one day after the final dose (day 17) or when they lost ≥15% of their body weight.

### Flow cytometry to measure cytokine concentrations

Peripheral blood was collected from the submandibular vein of 18-31-week-old mice into tubes containing lithium heparin (Sarstedt, Cat#41.1393.105) on day 3 at 18 h after the second poly(I:C) exposure of the chronic exposure timeline described above. Plasma was isolated by centrifuging at 700 xg for 15 mins at RT; plasma was then pulled off and placed into another tube that was centrifuged again at 2200 xg for 15 mins at RT then frozen at -80°C. Upon first freeze/thaw of plasma, cytokine protein levels were measured using the LEGENDplex Mouse Anti-Virus Response Panel (BioLegend, Cat# 740621) per manufacturer’s instructions as previously described (*21*). The LEGENDplex assay was evaluated on an Accuri C6 flow-cytometer then analyzed using LEGENDplex data analysis software. All samples were analyzed in duplicate and the average used for statistical analysis. Missing values were set to the lower limit of detection and values are only shown for cytokines detected above background.

### Embryo harvests

Male Dp16 were crossed overnight with 8-12-week-old synced female WT^1xIFNRs^. Dams were checked daily for vaginal plugs; the first morning of visual confirmation was denoted embryonic day (E)0.5. Upon 4-chamber heart formation at E15.5 (*80*), mouse embryos were harvested from dam after CO_2_ asphyxiation and cervical dislocation then embryos were briefly allowed to exsanguinate on ice in 1x PBS. A tail snip was collected from each embryo for genotyping.

### Histology to detect heart malformations

Upon harvest, embryos were fixed at 4°C overnight in 2% PFA (Ted Pella Inc, Cat#8505) diluted in 1x PBS then stored in 70% ethanol before embedding in paraffin. Embedded embryos were sectioned transversely at 7 µm thickness by using a LEICA RM 2155 Rotary Microtome. Serial sections were collected then hematoxylin and eosin Y (H&E) staining performed, followed by imaging with a Keyence BZ-X710 All-in-One Fluorescence Microscope. The investigators who sectioned embryos and performed H&E analysis were blind to embryo genotype.

### Sequencing and analysis of RNA from embryo hearts

Hearts were manually dissected from embryonic mice at E15.5 and RNA purified using Trizol reagent (manufacturer). RNA quality was assessed using an Agilent 2200 TapeStation and quantified on a Qubit fluorometer. Library preparation was carried out using a Universal Plus mRNA Kit Poly(A) (Tecan/Nugen). Paired-end, 150 bp sequencing was carried out on an Illumina NovaSeq 6000 instrument by the Genomics Shared Resource at the University of Colorado Anschutz Medical Campus. Subsequent analysis was carried out as for human Whole Blood RNA-seq, except alignment and gene-level count summarization used a mouse GRCm38 reference genome index and Gencode M24 annotation GTF. RNA-seq data yield was ∼84-143 × 10^6^ raw reads and ∼71-118 × 10^6^ final mapped reads per sample. Mouse embryonic heart RNA-seq data were deposited in the Gene Expression Omnibus GEO.

### Western blot of embryo hearts

Hearts were harvested from fresh embryos at E15.5 then frozen in liquid nitrogen. Frozen tissues were lysed in ice-cold lysis buffer (150 mM NaCl, 50 mM Tris-Cl pH 7.4, 1 mM EDTA, 1% Triton, with Complete mini tablet (Roche), 1 mM phenylmethylsulphonyl fluoride and 1x Halt™ Phosphatase Inhibitor Cocktail (Thermo Scientific). Protein concentration was determined by BCA assay and 10 µg protein lysates were loaded and resolved on a 10% polyacrylamide gel and transferred to a PVDF membrane (110mA, 16 hours, 4 °C). The following primary antibodies were used: total STAT1 (CST Cat# 9172; RRID:AB_2198300 1:1000), phospho-STAT1 (Tyr701) (CST Cat# 9167S; RRID:AB_561284 1:1000), and GAPDH (Thermo, Cat# AM4300; RRID:AB_2536381 1:5,000). Secondary antibodies used include the following: Goat Anti-Mouse IgG (Southern Biotech, Cat# 1031-05, 1:2,000) and Goat Anti-Rabbit IgG (Life Technologies, Cat# 65-6120, 1:2,000).

### Developmental milestones in neonates

Timeline for neonatal achievement of developmental milestones was assessed and analyzed as previously described (*81–85*) from days (D)3-21 post birth at 800-1100hr while experimenters were blind to genotype. Briefly, pups were removed from their home cage and placed in a holding cage with bedding maintained at 37°C by heating pad. Animals were identified by marking footpads or tattoos. Pups were assessed in a pseudorandom order by blinded investigators for developmental milestone criteria in the following order: 1) surface righting 2 days in a row (D3-10), a righting reflex that determines ability of the pup for body-righting when placed upside down; 2) first day both eyes open (D7-21); 3) first day both ears twitch (D7-21), a touch sensitivity test; and 4) the first day of auditory startle (D11-21), a test of sensitivity to a sharp loud click noise. Once criteria were reached, testing was stopped for that mouse. The investigators who assessed developmental milestones were blind to neonate genotype.

To test for differences in the chance of success in achieving each developmental milestone, results were treated as time-to-event data and analyzed using a mixed effects Cox regression approach using the survival (version 3.2-7 (*86*)), coxme (version 2.2-16 (*87*)), emmeans (version 1.5.1 (*88*),) and broom (version 0.7.9 (*89*)) packages in in R. Models for each milestone were generated using the *coxme()* function from the coxme package with a time-to-event survival object as the outcome variable, genotype as the variable of interest, and with adjustment for sex and sex*genotype interaction as fixed effects and litter as a random effect. Hazard ratios (referred to herein as “Success ratios”), 95% confidence intervals (CIs), and p-values for all pair-wise genotype comparisons, either combined or stratified by sex, were obtained from model objects using the *emmeans()* and *contrast()* functions from the emmeans package, and FDR q-values calculated using the Benjamini-Hochberg method. Data are presented in **Fig. 4A** and **Fig. S5A** as success ratios with size proportional to q-value and 95% CIs for each pair-wise genotype comparison.

### Cognitive tests in adulthood

Mice of 4-5 months of age were handled two mins per day for 2-5 days leading up to the first test. When mice were run through multiple assays, they were run through rotarod, Morris water maze (MWM), then contextual fear conditioning (CFC). For all assays, mice were allowed to acclimate to the experimental room in their home cages for 30 mins before testing began each day at 1200-1600 h. Equipment was cleaned in between mice.

The rotarod performance test was used to measure motor coordination as previously described for mouse models of DS (*90*). Briefly, mice were placed on a rotating cylinder at a set speed and had to keep pace with the rotation to stay on the cylinder (*28*). Each mouse was given 2 practice sessions at 16 revolutions per minute (RPM), followed by 2 test sessions at 16, 24, and 32 RPM. Each session ended after 120 s or when the mouse fell. Inter-session intervals were 15 mins. Latency to fall was tracked using Rotarod Version1.4.1 (Copyright 2002-2010, MED Associates, Inc.) and averaged across both test sessions for each speed.

MWM was performed on mice to assess spatial learning and memory as previously described for preclinical models of DS with slight modifications (*90–92*). The MWM consisted of a pool 120 cm in diameter filled with opaque water in which an escape platform was hidden at 30 cm away from the center of the area; mice must learn then remember extra-maze visual cues from around the room to efficiently navigate to the submerged platform during successive swim trials (**Fig. S5C**).

Visible platform trials were not conducted as it was previously shown that Dp16 and WT mice show no differences (*90, 93, 94*), although this was not always the case (*82*). Instead, mice started with one unrecorded swim trial where they were directed to the platform by a visual cue on the platform. Mice were then released into the MWM at pseudorandomized locations of the pool edge (i.e., N, S, E, and W locations of **Fig. S5C-D**) for a total of four swim trials per MWM block and two blocks per day with the final reported experimental value per block reflecting the average of the four swim trials. Each swim trial concluded when the mouse found the hidden platform or after 1 min, whichever came first. Immediately following block 6, mice underwent a 1 min probe trial where the platform was removed to conclude the acquisition phase of swim trials 1-24. After the first probe, we immediately employed a reversal phase for another 6 blocks of swim trials 25-48 that immediately precluded a second 1 min Probe Trial. During the Acquisition Phase, mice learn the location of the platform while in the Reversal Phase mice must relearn to find the platform again after it is moved to the opposite quadrant of the pool (**Fig. S5D**). Throughout both phases, more efficient navigation to the platform was interpreted as superior allocentric memory of the platform location (**Fig. S5C**).

Swim data were collected using the video tracking system Ethovision (Noldus) v8.5. Nesting was applied to swim paths in Ethovision prior to analysis where the center point (mouse) must be between start and stop threshold velocities of 2 and 1.75 cm/s to avoid giving weight of initial interaction of animal to arena and tracking of experimenters’ hand during mouse drop into the maze.

The CFC test of associative learning and memory was performed as previously described for mouse models of DS (*90*). Briefly, CFC boxes (30.5×24.1×29.2 cm) consisted of a light, recording device, and had a metal grid on the floor (Med Associates, St. Albans, VT, Modular Mouse Test Chamber). During the 5 min training phase, mice explored the enclosure freely before receiving two 0.5 mA foot shocks (2 s) at 180 s and 240 s. Testing phase occurred exactly 24 h after training phase and was identical to training without foot shocks where mice were exposed to the same context to test for memory of previous shocks. During both phases, freezing behavior was measured FreezeScan version 2.00 (Copyright 2000-2002, Clever Sys. Inc.) and interpreted as fear in the testing phase.

### Morphometric analysis skull form

Mice were decapitated at 7-8 weeks of age then whole heads imaged using a SkyScan 1275 (μCT). Scanning was performed at 17.6-micron resolution using the following parameters: 55 kV, 180 mA, 0.5 mm Al filter; 0.3° rotation step over 180°, and 3-frame averaging. All raw scan data were reconstructed to multiplanar slice data using NRecon V1.7.4.6 software (Bruker, Belgium). Reconstructed data were then rendered in 3D with consistent thresholding parameters using Drishti V2.6.5 Volume Exploration software (Limaye, 2012) for gross visual assessment of the craniofacial skeleton. Representative rendered images were captured and processed using Photoshop (Adobe Creative Cloud).

Morphometric analysis of craniofacial landmarks (LMs) was then used to compare skull form (e.g. skull size and shape) between genotypes as previously described (*30*). Briefly, coordinates for 30 homologous LMs were independently collected from each 3D rendered skull within Drishti V2.6.5 by two investigators blinded to genotype (**Fig. S6A** and **Data S6**). LM datasets from two investigators were averaged and then normalized by their respective root centroid size (RCS) values to remove overall skull size as a variable and are reported in voxel units (**Data S6**). The WinEDMA package was used to conduct Euclidean Distance Matrix Analysis (EDMA), which analyzes morphological differences between two groups of specimens by assessing the change in ratio values between respective LM pairs (*31*). Following Lele and Richtsmeier (*95*), the 90% CIs were calculated by bootstrapping the shape difference matrix 10,000 times (**Data S6**). The FORM procedure was employed to find inter-LM distances that differed between populations, as well as LMs influencing or driving those differences (*95*). Inter-LM distances were deemed different between populations if the CIs did not cross 1.

### Qualitative assessment of the skull

Except for calvaria rounding, which was done on mice 7-10 weeks of age, all qualitative assessment of skulls from decapitated mice were done at 7-8 weeks of age by an expert in craniofacial morphology blind to genotypes. As mineralization of spheno-occipital synchondrosis (SOS) typically begins around postnatal D28 to ultimately mediate fusion of the basisphenoid (BS) and basioccipital bones by 12 weeks of age (*30*), unusually pronounced ossification of the intersphenoid synchondrosis (ISS) was formally assessed. An ISS severity score was assigned using the following criteria: 0 = normal unfused appearance; 1 = <1/3^rd^ of the ISS width bridged by ossification; 2 = between 1/3^rd^ and 2/3^rd^ of the ISS bridged by ossification; 3 = the presphenoid and BS bones of the anterior cranial base were completely fused, obliterating the ISS.

### Statistical analysis and result visualization

Sample size was determined a priori based on effect sizes of previous studies (e.g., human studies) or by post hoc analyses to ensure >80% power was achieved to reduce type II error (e.g., heart histology, developmental milestone achievement, and cognition in adults). All statistical analyses were conducted in GraphPad Prism version 8.0.1 (RRID: SCR_002798) or R and are listed with sample sizes in the corresponding figure legends and described above. Preprocessing, statistical analysis, and initial plot generation for all datasets was carried out using R (R 4.0.1; RRID:SCR_001905) (R Studio; RRID:SCR_000432) (*96, 97*). Method schematics were generated using Biorender.com. All figures were finalized in Adobe Illustrator V24.1.

---

**Fig. S1.**
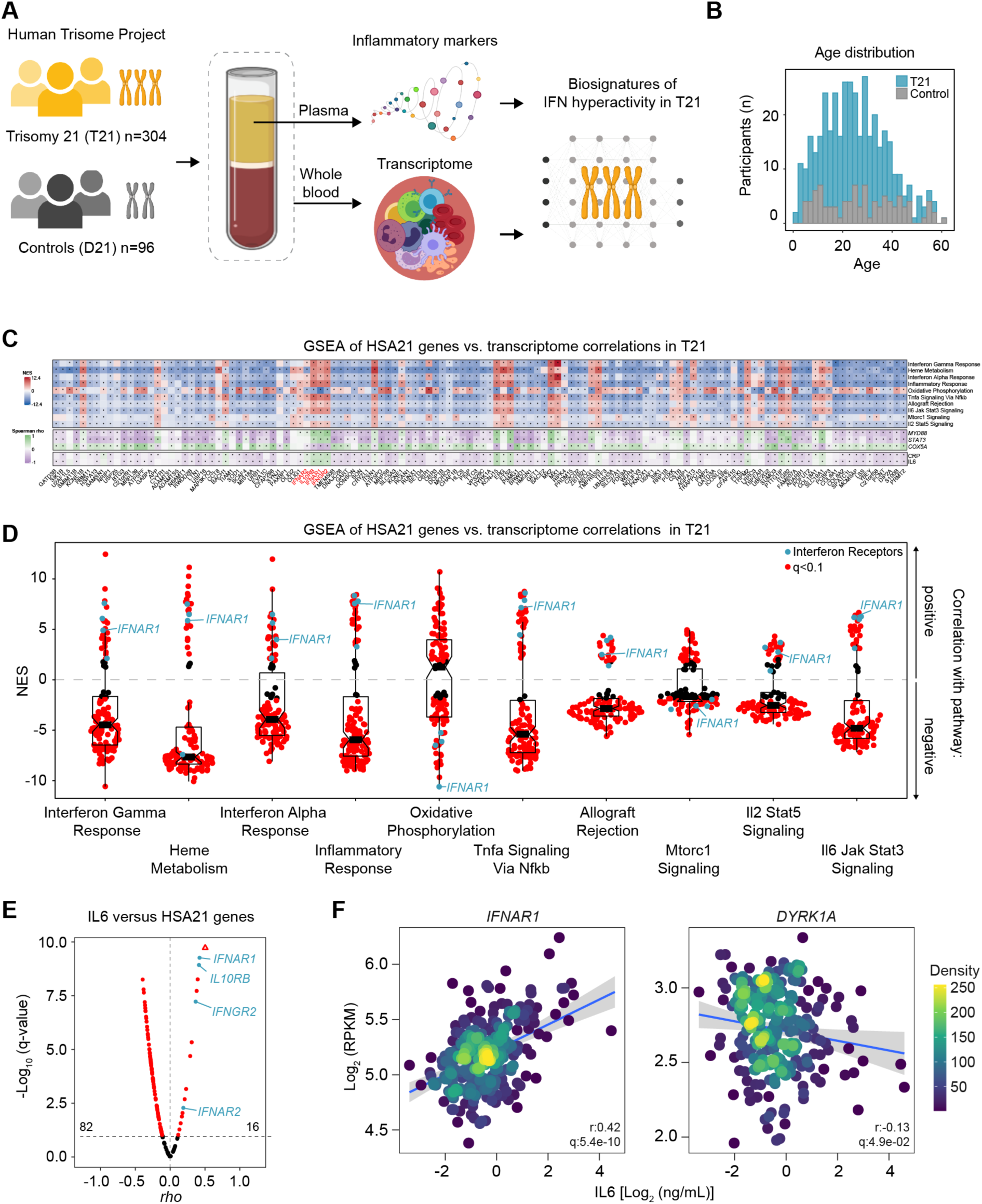
Overexpression of *IFNRs* associates with inflammation in people with trisomy 21. **(A)** Schematic of biospecimen source and processing for all datasets underlying Fig. 1 and Fig. S1. **(B)** Age distribution of cohorts analyzed. **(C) Top:** Heatmap displaying the results of gene set enrichment analysis (GSEA) of Spearman *rho* value correlations between transcripts encoded on human chromosome 21 (HSA21, y-axis) versus the rest of the transcriptome, using only samples from individuals with T21. Only the top 10 gene pathways activated in the transcriptome of individuals with Down syndrome are shown. NES: normalized enrichment score. **Middle:** Spearman correlations between all transcripts encoded on HSA21 and the indicated differentially expressed genes encoded elsewhere in the genome among individuals with T21. **Bottom:** Spearman correlations between transcripts encoded on HSA21 and the indicated inflammatory proteins measured in plasma among individuals with T21. *q<0.1 **(D)** Sina plots displaying the distribution of NES values after running GSEA on Spearman *rho* values for the correlation between expression of transcripts encoded on HSA21 and the top 10 gene signatures elevated in the transcriptome of individuals with Down syndrome. Only T21 samples were used for this analysis. **(E)** Volcano plot of *rho* and q-values from Spearman correlations of IL6 protein abundance in plasma versus expression of HSA21 genes in the whole blood transcriptome of individuals with T21. **(F)** XY scatter plots show Spearman correlations between the indicated mRNAs and IL6 protein abundance. In (E-F, statistical significance was determined by Benjamini-Hochberg correction of Spearman p values and a threshold of 0.1 (*q<0.1).

**Fig. S2.**
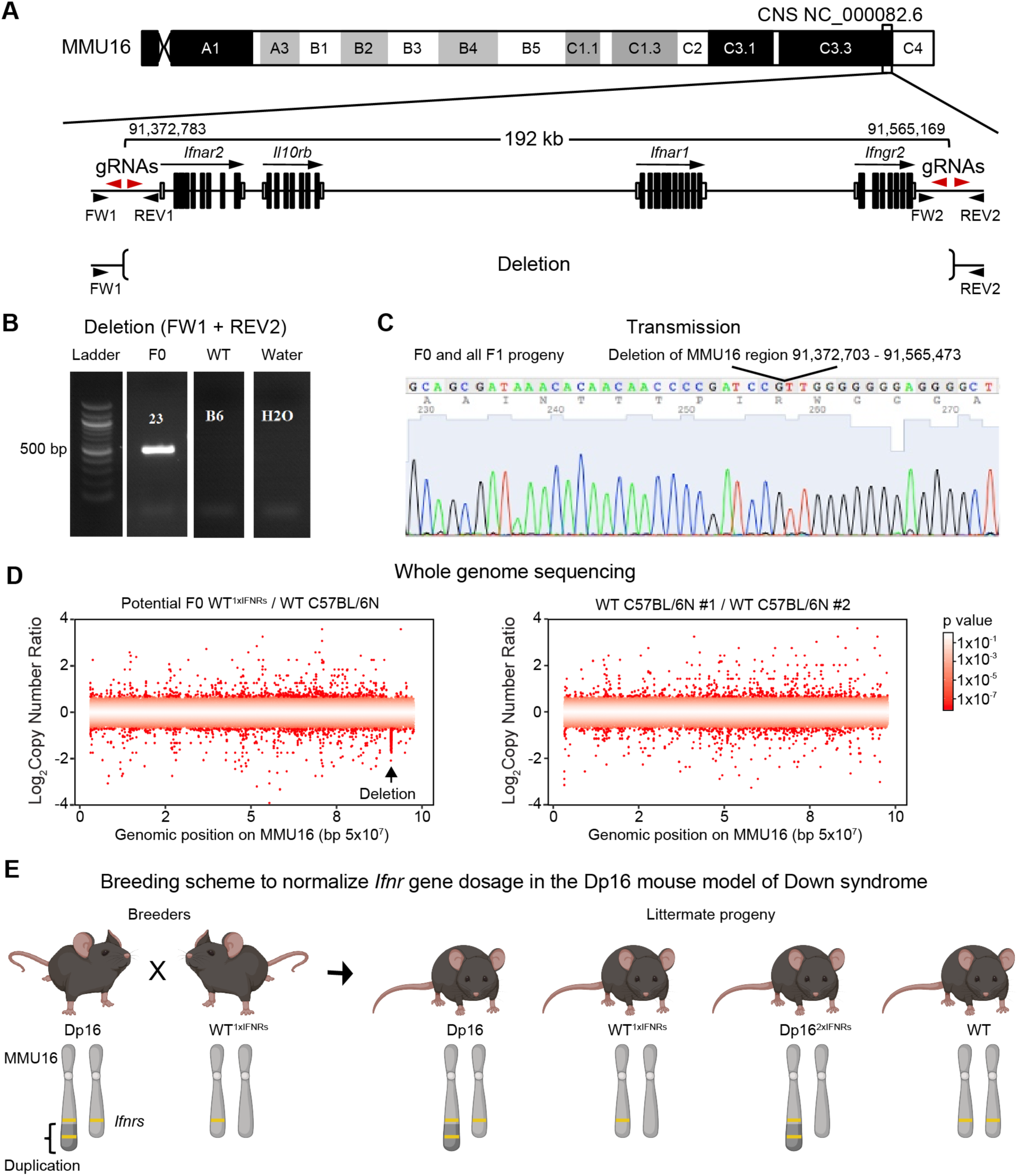
Novel mouse model to determine if triplication of *Ifnr* genes is necessary for phenotypes of Down syndrome. **(A)** Schematic of the murine interferon receptor (*Ifnr)* gene locus and CRIPSR/Cas9 guide RNAs (gRNAs, red arrowheads) used for the deletion of a 192 kb genomic region of chromosome 16 (MMU16). Forward and reverse genotyping primers (FW and REV, black arrowheads) are indicated. Molecular location of genes based on GRCm38 reference genome is indicated on an ideogram of MMU16 cytogenetic regions colored according to Giemsa banding. **(B)** PCR of DNA from the founder (F0) mouse bearing the expected deletion (+) and negative controls [-; wildtype (WT) and Water (no DNA)]. **(C)** Representative Sanger sequencing of the single modified allele transmitted from the F0 male to the first generation of progeny (F1) after intercrossing with a WT female. **(D)** Whole genome sequencing followed by copy number variant analysis from the F0 mouse bearing a deletion of the *Ifnr* locus syntenic to human chromosome 21 (WT^1xIFNRs^) with the site of deletion on MMU16 denoted by arrow (left image) that is absent when two non-related C57BL/6N WT mice are compared (right image). *p<0.1 by CNV-seq. **(E)** Diagram of breeding strategy to reduce copy number of just four *Ifnr* genes (yellow line) triplicated in the Dp16 mouse model of Down syndrome by intercrossing Dp16 males with WT^1xIFNRs^ females. Littermate progeny can then be compared between Dp16, Dp16 normalized for just *Ifnr* dose (Dp16^2xIFNRs^), and WT controls.

**Fig. S3.**
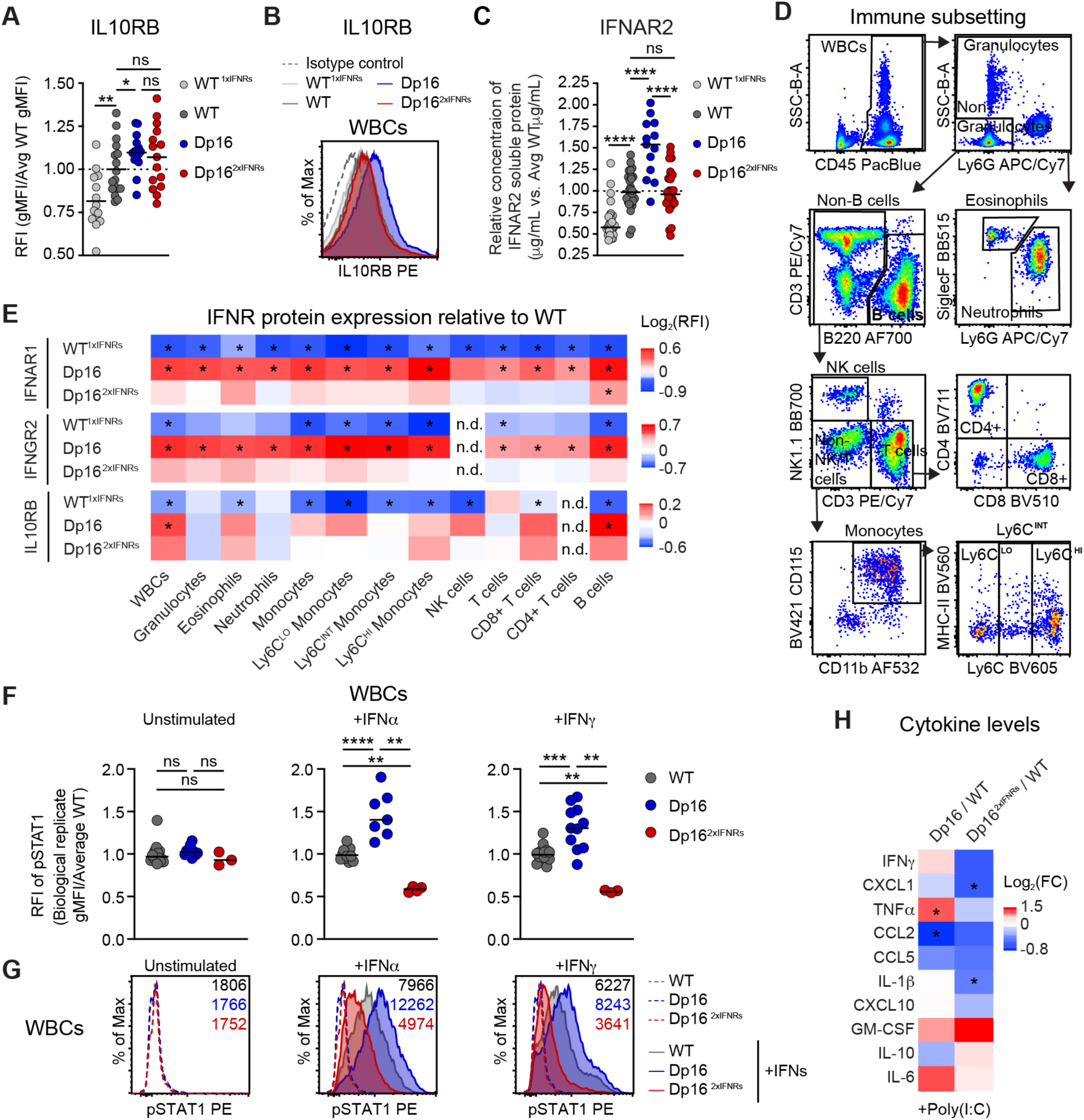
Triplication of *Ifnr* genes drives increased IFNR protein expression and an aberrant antiviral response in a mouse model of trisomy 21. **(A)** Relative fluorescent intensity (RFI) of the geometric mean fluorescent intensity (gMFI) and **(B)** example histogram for IL10RB protein on the surface of CD45+ white blood cells (WBCs) from heterozygous *Ifnr* knockout mice, (WT^1xIFNRs^), wild type mice (WT), Dp16 (Down syndrome) mice, and Dp16 normalized for just *Ifnr* gene dosage (Dp16^2xIFNRs^), as measured by flow cytometry (n=14-17/group). **(C)** Soluble IFNAR2 protein detected by ELISA in the blood of each biological replicate relative to the average WT value per experiment (n=13-28/group). (**D**) Gating strategy of immune subsets in whole blood by flow cytometry. (**E**) IFNR protein expression on immune subsets from (D) (n=5-28/group). Heatmap indicates RFIs for gMFIs of the indicated genotype over the WT average IFNR gMFI per experiment. n.d. denotes IFNR RFIs that were not detected above isotype background. **(F-G)** Phosphorylation of STAT1 (pSTAT1) was analyzed via flow cytometry for CD45+ WBCs at baseline or after stimulation with IFNα (10,000 U/mL) or IFNγ (100 U/mL) for 30 minutes. Data for each genotype are presented as RFIs for gMFIs relative to the average WT value per experiment (F) or example histograms with gMFIs indicated (G). **(H)** Cytokine levels in the plasma of mice treated with poly(I:C). Heatmap indicates Log_2_fold-change of cytokine protein in plasma of the indicated cohort + poly(I:C) relative to the WT + poly(I:C) cohort after 3 days of the poly(I:C) regimen. In A, C and F, each dot represents an independent biological replicate with the mean indicated. Significance by Mann Whitney test denoted by *p≤0.05, **p≤0.01, ***p≤0.001, and ****p≤0.001.

**Fig. S4.**
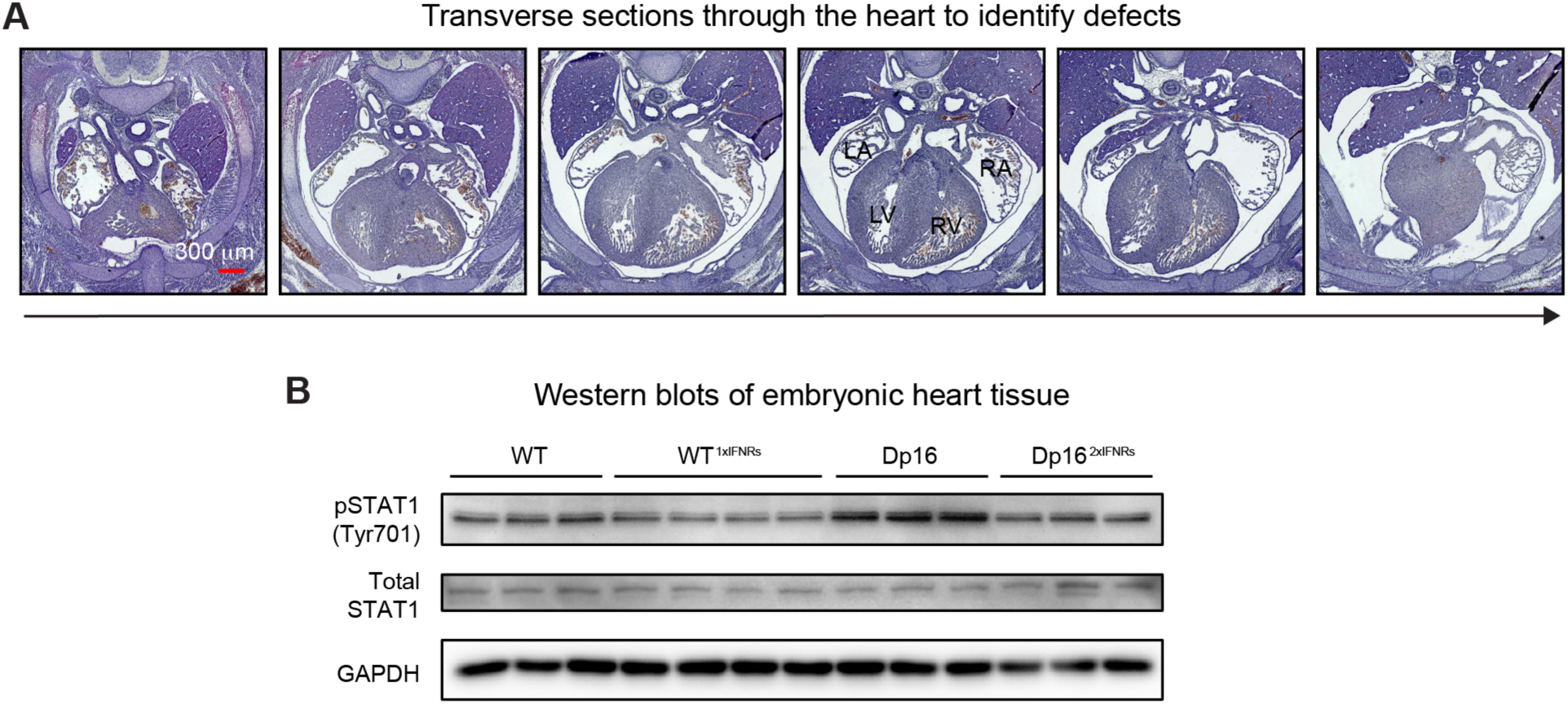
Normalization of *Ifnr* copy number prevents embryonic heart malformations in a mouse model of DS. **(A)** Representative hematoxylin and eosin-stained sections of a heart in murine embryos at embryonic day (E)15.5. Serial sections were cut through the entire region of the developing murine heart. **(B)** Western blot analysis of total and phosphorylated STAT1 protein from developing hearts at E15.5 from heterozygous *Ifnr* knockout mice, (WT^1xIFNRs^), wildtype mice (WT), Dp16 (Down syndrome) mice, and Dp16 normalized for just *Ifnr* gene dosage (Dp16^2xIFNRs^) (n=3-4/group).

**Fig. S5.**
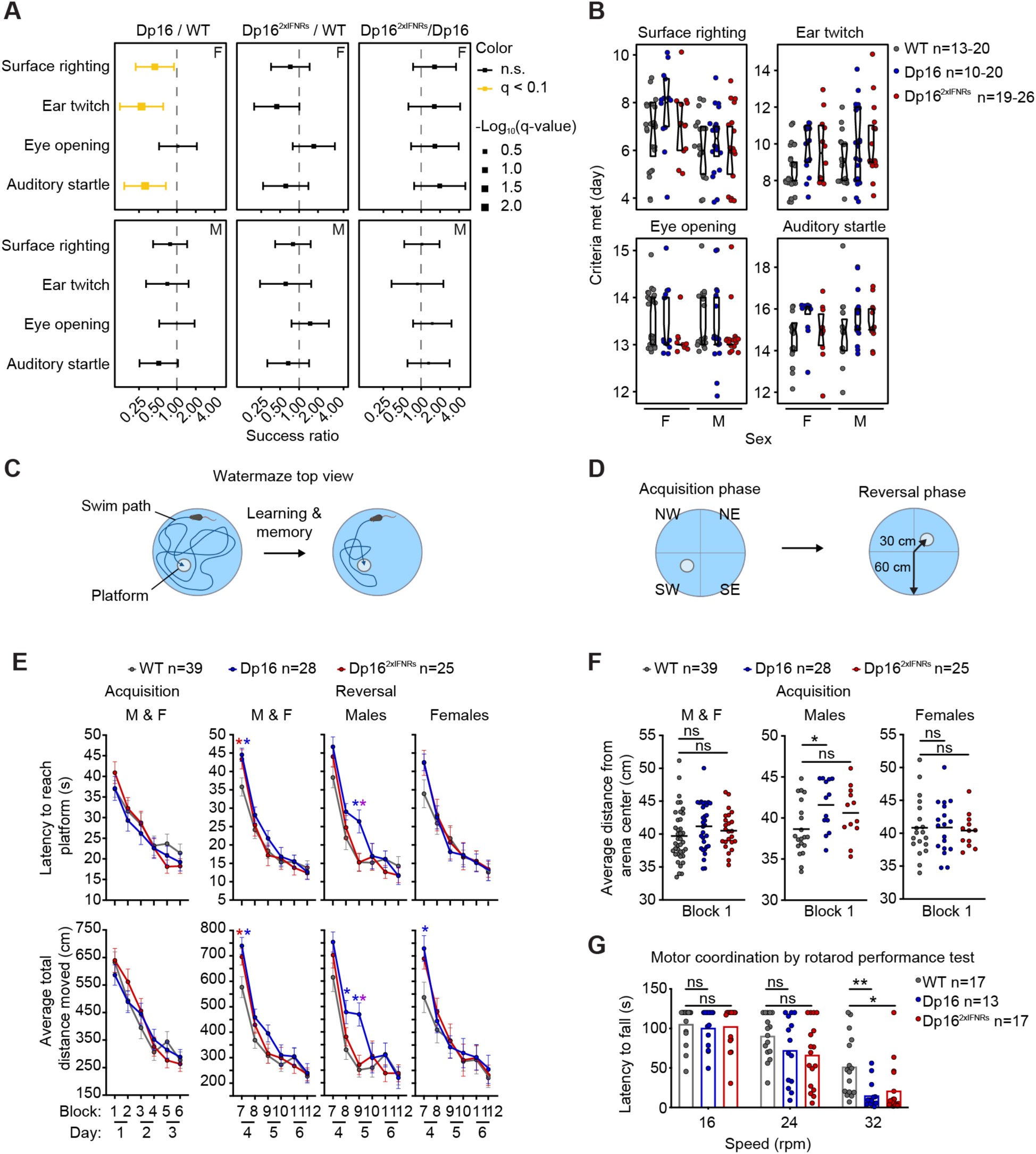
Cognitive and behavioral deficits are ameliorated by normalization of *Ifnr* copy number in a mouse model of trisomy 21. **(A)** Success ratio of developmental milestone achievement in neonates 3-21 days old by Cox regression after adjusting for gender and litter with Benjamini-Hochberg correction. Averages for females (F) and males (M) are flanked by confidence intervals; cohorts represent wildtype (WT) controls (n=43-46, 19-20 male), the Dp16 preclinical mouse model of Down syndrome (n=32-34, 19-20 male), and Dp16 normalized for gene dose of just the *Ifnrs* (Dp16^2xIFNRs^; n=27-30, 16-20 male). (**B**) Day of developmental milestone achievement by genotype. Raw unadjusted data are presented as modified sina plots with boxes representing medians and upper/lower quartiles (n=10-26/group). **(C-D)** Schematic of Morris water maze (MWM). With learning and memory, the mice use visual cues to navigate more efficiently to a hidden platform (C). Swims are divided among two successive phases where the platform is located in opposite quadrants that are labeled by intercardinal directions (D). **(E)** Duration (top) and distance traveled (bottom) until platform escape during swims in the MWM for all male and female (M & F) WT controls (n=39, 19 male), Dp16 (n=28, 13 male), and Dp16^2xIFNRs^ (n=25, 12 male). Data represent the average +/- SEM for mice 4-5 months in age with statistical significance at p≤0.05 by two-way repeated measures ANOVA with post-hoc Tukey’s HSD. Significant differences between pairs are denoted by a blue * for Dp16 versus WT, a red * for Dp16^2xIFNRs^ versus WT, and a purple * for Dp16 versus Dp16^2xIFNRs^. **(F)** Tendency of mice to swim near the periphery of the MWM environment in the first block. **(G)** Performance in rotarod measured by time until fall from a rod rotating at the indicated speed (n=13-17/group, 6-8 male). In F-G scatter and bar graphs represent the average +/- range indicated by individual biological replicates(circles) of mice 4-5 months of age where *p≤0.05 and **p≤0.01 by one-way ANOVA with post-hoc Tukey’s HSD.

**Fig. S6.**
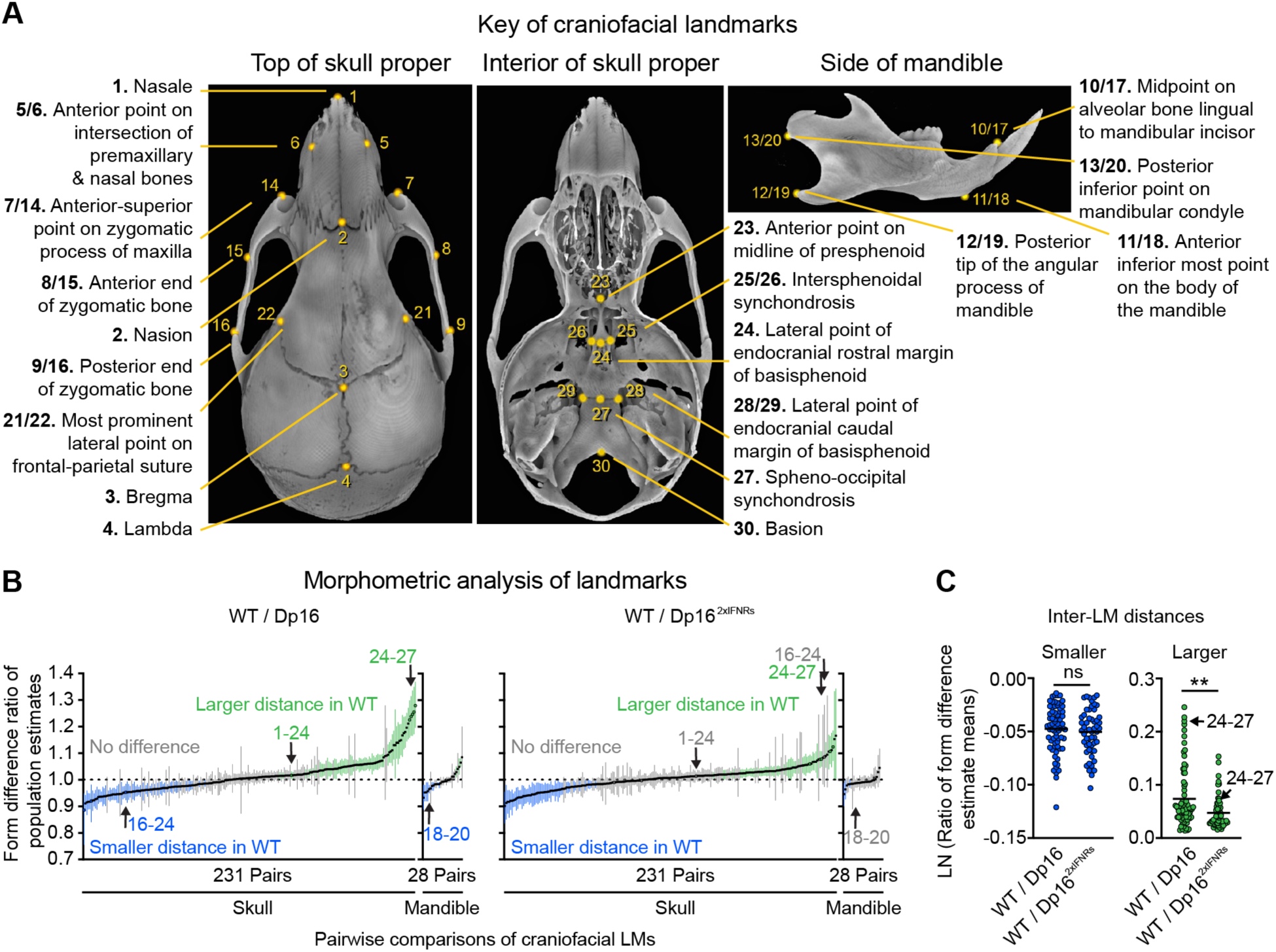
Triplication of *Ifnr* genes exacerbates craniofacial morphometric differences in a mouse model of Down syndrome. **(A)** Representative micro-computed tomography (μCT) images of a wildtype (WT) mouse skull. Single and paired landmarks (LMs) are denoted on dorsal views of the outer skull proper (left image) and interior cranial base (center image) as well as on a lateral view of the hemi-mandible (right image). **(B)** Form difference ratio of inter-LM distances of the skull proper and mandible by Euclidean Distance Matrix Analysis (EDMA) for cohorts of Dp16 (Down syndrome) or Dp16 normalized for just *Ifnr* gene dosage (Dp16^2xIFNRs^), compared to WT controls (n=6-7/group followed by bootstrapping 10,000x). Mean population estimates (circles) are on confidence intervals (CIs, lines) and colored according to differences where green represents larger distance for the numerator (i.e. CIs>1), blue represents smaller distances for the numerator (i.e. CIs<1), and black/grey represents no change in distances (i.e. CIs that intersect 1). **(C)** Natural Log (LN) of form difference ratio of all mean population estimates for inter-LM distances of the skull proper and mandible shown in panel (B) that were smaller (blue) or larger (green) in WT animals. Significance was determined by Mann-Whitney test where *p≤0.05 and **p≤0.01.

**Fig. S7.**
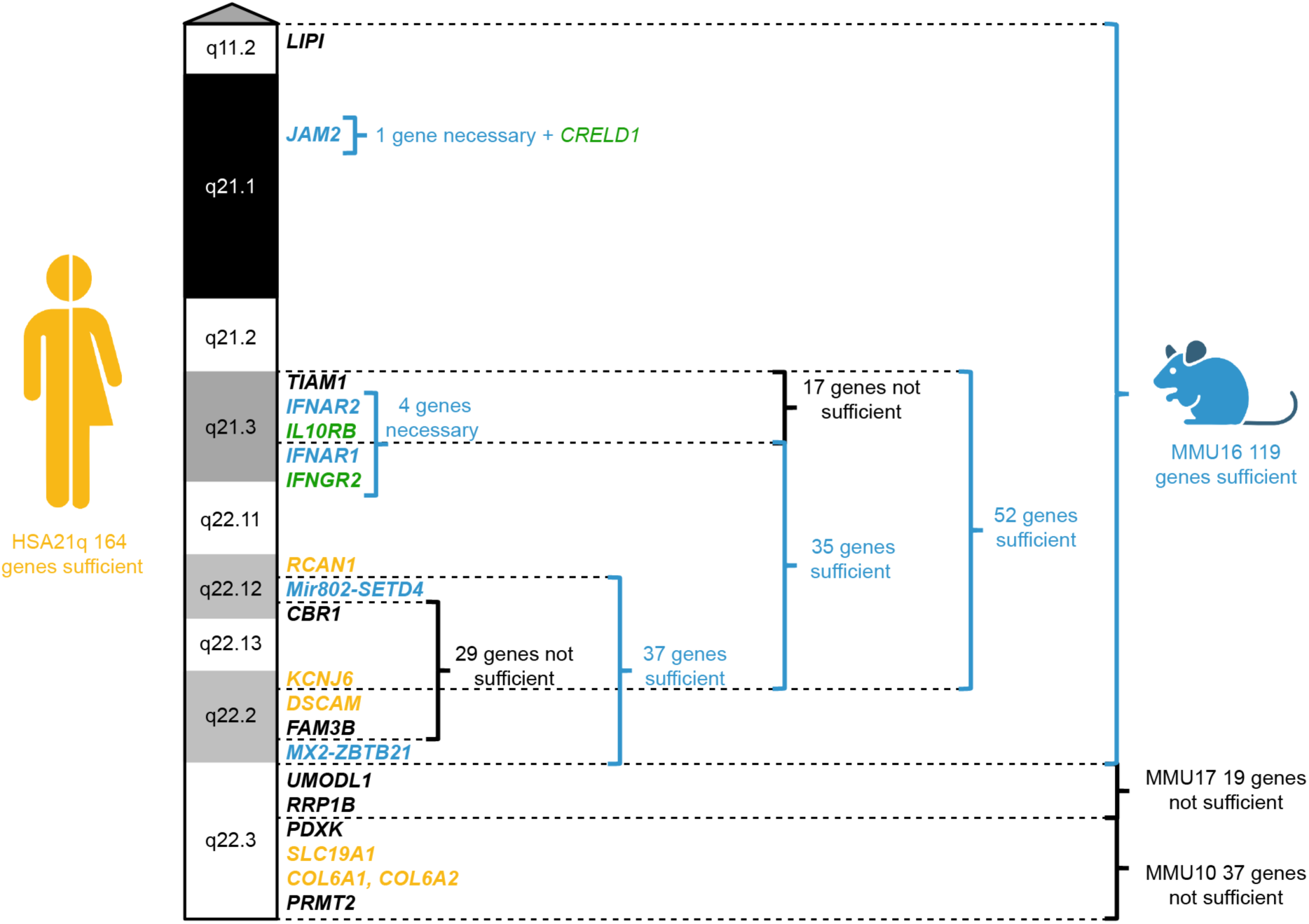
Overview of experimental evidence regarding the genetic basis of heart defects in mouse models of Down syndrome. Diagram depicts genetic variants on human chromosome 21 (HSA21) that may contribute to risk of congenital heart defects (CHDs) in humans with trisomy 21 (T21, yellow), genes with functional evidence whose triplication is necessary or sufficient to increase incidence of CHDs in mouse models of Down syndrome (DS, blue), and genes with supporting evidence in both humans and mouse models of DS (green). Relative cytogenetic locations and number of protein-coding genes (bold) are indicated along ideogram of the q arm for HSA21 colored according to Giemsa banding (*3, 16, 23, 24, 56-58*).

**Data S1. Demographics of human research cohort (separate file).**

**Data S2. Whole blood transcriptome analysis (separate file).**

Tab A. RNA sequencing (RNA-seq) results of whole blood compared between people with T21 and typical D21 controls.

Tab B. Gene Set Enrichment Analysis (GSEA) of whole blood transcriptome after comparison between people with T21 and typical D21 controls.

Tab C. GSEA of HSA21 transcripts from whole blood transcriptome after comparison between people with T21 and typical D21 controls.

Tab D. Spearman correlation of *IFNAR1* transcript abundance by the entire whole blood transcriptome from people with T21.

Tab E. Spearman correlation of *DYRK1A* transcript abundance by the entire whole blood transcriptome from people with T21.

Tab F. Spearman correlation of *MYD88* transcript abundance by the entire whole blood transcriptome from people with T21.

Tab G. Spearman correlation of *STAT3* transcript abundance by the entire whole blood transcriptome from people with T21.

Tab H. Spearman correlation of *COX5A* transcript abundance by the entire whole blood transcriptome from people with T21.

**Data S3. Plasma inflammatory marker analysis (separate file).**

Tab A. Mean concentrations of CRP from the plasma of individuals with T21 samples in duplicate.

Tab B. Mean concentrations of IL6 from the plasma of individuals with T21 samples in duplicate.

Tab C. Spearman correlation of CRP protein abundance versus HSA21 encoded transcripts measured in the whole blood transcriptome from people with T21.

Tab D. Spearman correlation of IL6 protein abundance versus HSA21 encoded transcripts measured in the whole blood transcriptome from people with T21.

**Data S4. Additional method data (separate file).**

Tab A. Sequences of guide RNAs (gRNAs) that flank interferon receptor (*Ifnr*) genes encoded on mouse chromosome 16 (MMU16).

Tab B. PCR strategy to screen mutant mice for large chromosomal rearrangements on MMU16.

Tab C. PCR strategy for genotyping of mice used in this study.

Tab D. qRT-PCR strategy to measure *Ifnrs* expression.

**Data S5. Transcriptome analysis of embryonic hearts (separate file).**

Tab A. RNA-seq results of E15.5 hearts compared between Dp16 and WT mice.

Tab B. RNA-seq results of E15.5 hearts compared between Dp16^2xIFNRs^ and WT mice.

Tab C. RNA-seq results of E15.5 hearts compared between Dp16^2xIFNRs^ and Dp16 mice.

Tab D. GSEA results of the E15.5 heart transcriptome after comparison between Dp16 and WT mice.

Tab E. GSEA results of the E15.5 heart transcriptome after comparison between Dp16^2xIFNRs^ and Dp16 mice.

Tab F. GSEA results of the E15.5 heart transcriptome after comparison between Dp16^2xIFNRs^ and WT mice.

**Data S6. Craniofacial analysis by micro-computed tomography (separate file).**

Tab A. Raw inter-landmark (LM) distances.

Tab B. Inter-LM distances from Tab A after normalization of each skull to their root centroid size (RCS).

Tab C. Form-difference ratio estimates of population minimum, mean, and maximum values after bootstrapping of inter-LM distances in Tab B.

## References and Notes

1. J. Lejeune, R. Turpin, M. Gautier, [Mongolism; a chromosomal disease (trisomy)]. Bull Acad Natl Med 143, 256–265 (1959).

2. C. T. Mai et al., National population-based estimates for major birth defects, 2010-2014. Birth Defects Res 111, 1420–1435 (2019).

3. S. E. Antonarakis, et al., Down syndrome. Nature reviews. Disease primers 6, 9 (2020).

4. B. Chicoine et al., Prevalence of Common Disease Conditions in a Large Cohort of Individuals With Down Syndrome in the United States. J Patient Cent Res Rev 8, 86–97 (2021).

5. A. K. Clift, C. A. C. Coupland, R. H. Keogh, H. Hemingway, J. Hippisley-Cox, COVID-19 Mortality Risk in Down Syndrome: Results From a Cohort Study Of 8 Million Adults. Ann Intern Med, (2020).

6. K. D. Sullivan et al., Trisomy 21 consistently activates the interferon response. Elife 5, (2016).

7. J. W. Schoggins, Interferon-Stimulated Genes: What Do They All Do? Annu Rev Virol 6, 567–584 (2019).

8. M. Hattori et al., The DNA sequence of human chromosome 21. Nature 405, 311–319 (2000).

9. Y. H. Tan, E. L. Schneider, J. Tischfield, C. J. Epstein, F. H. Ruddle, Human chromosome 21 dosage: effect on the expression of the interferon induced antiviral state. Science 186, 61–63 (1974).

10. R. K. Powers et al., Trisomy 21 activates the kynurenine pathway via increased dosage of interferon receptors. Nat Commun 10, 4766 (2019).

11. K. A. Waugh et al., Mass Cytometry Reveals Global Immune Remodeling with Multi-lineage Hypersensitivity to Type I Interferon in Down Syndrome. Cell reports 29, 1893–1908 e1894 (2019).

12. M. Krivega et al., Genotoxic stress in constitutive trisomies induces autophagy and the innate immune response via the cGAS-STING pathway. Commun Biol 4, 831 (2021).

13. M. P. Rodero, Y. J. Crow, Type I interferon-mediated monogenic autoinflammation: The type I interferonopathies, a conceptual overview. The Journal of experimental medicine 213, 2527–2538 (2016).

14. L. Malle, D. Bogunovic, Down syndrome and type I interferon: not so simple. Curr Opin Immunol 72, 196–205 (2021).

15. K. J. Gardiner, Pharmacological approaches to improving cognitive function in Down syndrome: current status and considerations. Drug Des Devel Ther 9, 103–125 (2015).

16. Z. Li et al., Duplication of the entire 22.9 Mb human chromosome 21 syntenic region on mouse chromosome 16 causes cardiovascular and gastrointestinal abnormalities. Hum Mol Genet 16, 1359–1366 (2007).

17. J. M. Starbuck, T. Dutka, T. S. Ratliff, R. H. Reeves, J. T. Richtsmeier, Overlapping trisomies for human chromosome 21 orthologs produce similar effects on skull and brain morphology of Dp(16)1Yey and Ts65Dn mice. American journal of medical genetics. Part A 164A, 1981–1990 (2014).

18. J. W. Goodliffe et al., Absence of Prenatal Forebrain Defects in the Dp(16)1Yey/+ Mouse Model of Down Syndrome. The Journal of neuroscience : the official journal of the Society for Neuroscience 36, 2926–2944 (2016).

19. T. Yu et al., Effects of individual segmental trisomies of human chromosome 21 syntenic regions on hippocampal long-term potentiation and cognitive behaviors in mice. Brain research 1366, 162–171 (2010).

20. N. M. Aziz et al., Lifespan analysis of brain development, gene expression and behavioral phenotypes in the Ts1Cje, Ts65Dn and Dp(16)1/Yey mouse models of Down syndrome. Dis Model Mech 11, (2018).

21. K. D. Tuttle et al., JAK1 Inhibition Blocks Lethal Immune Hypersensitivity in a Mouse Model of Down Syndrome. Cell reports 33, 108407 (2020).

22. N. J. Espat, E. M. Copeland, L. L. Moldawer, Tumor necrosis factor and cachexia: a current perspective. Surg Oncol 3, 255–262 (1994).

23. C. Liu et al., Genetic analysis of Down syndrome-associated heart defects in mice. Hum Genet 130, 623–632 (2011).

24. C. Liu et al., Engineered chromosome-based genetic mapping establishes a 3.7 Mb critical genomic region for Down syndrome-associated heart defects in mice. Hum Genet 133, 743–753 (2014).

25. P. Curzon, N. R. Rustay, K. E. Browman, in Methods of Behavior Analysis in Neuroscience, nd, J. J. Buccafusco, Eds. (Boca Raton (FL), 2009).

26. R. Morris, Developments of a water-maze procedure for studying spatial learning in the rat. Journal of neuroscience methods 11, 47–60 (1984).

27. D. Treit, M. Fundytus, Thigmotaxis as a test for anxiolytic activity in rats. Pharmacol Biochem Behav 31, 959–962 (1988).

28. N. W. Dunham, T. S. Miya, A note on a simple apparatus for detecting neurological deficit in rats and mice. J Am Pharm Assoc Am Pharm Assoc 46, 208–209 (1957).

29. J. M. Starbuck, T. M. Cole, 3rd, R. H. Reeves, J. T. Richtsmeier, The Influence of trisomy 21 on facial form and variability. American journal of medical genetics. Part A 173, 2861–2872 (2017).

30. S. R. Vora, E. D. Camci, T. C. Cox, Postnatal Ontogeny of the Cranial Base and Craniofacial Skeleton in Male C57BL/6J Mice: A Reference Standard for Quantitative Analysis. Front Physiol 6, 417 (2015).

31. T. M. Cole, 3rd, J. T. Richtsmeier, A simple method for visualization of influential landmarks when using euclidean distance matrix analysis. Am J Phys Anthropol 107, 273–283 (1998).

32. J. J. Alio, J. Lorenzo, C. Iglesias, Cranial base growth in patients with Down syndrome: a longitudinal study. Am J Orthod Dentofacial Orthop 133, 729–737 (2008).

33. S. Rengasamy Venugopalan, E. Van Otterloo, The Skull’s Girder: A Brief Review of the Cranial Base. J Dev Biol 9, (2021).

34. L. E. Maroun, Interferon action and chromosome 21 trisomy (Down syndrome): 15 years later. Journal of theoretical biology 181, 41–46 (1996).

35. K. Boroviak, B. Doe, R. Banerjee, F. Yang, A. Bradley, Chromosome engineering in zygotes with CRISPR/Cas9. Genesis 54, 78–85 (2016).

36. X. F. Kong et al., Three Copies of Four Interferon Receptor Genes Underlie a Mild Type I Interferonopathy in Down Syndrome. J Clin Immunol 40, 807–819 (2020).

37. T. Illouz et al., Immune Dysregulation and the Increased Risk of Complications and Mortality Following Respiratory Tract Infections in Adults With Down Syndrome. Front Immunol 12, 621440 (2021).

38. A. K. Clift, C. A. C. Coupland, R. H. Keogh, H. Hemingway, J. Hippisley-Cox, COVID-19 Mortality Risk in Down Syndrome: Results From a Cohort Study of 8 Million Adults. Ann Intern Med 174, 572–576 (2021).

39. A. Huls et al., Medical vulnerability of individuals with Down syndrome to severe COVID-19-data from the Trisomy 21 Research Society and the UK ISARIC4C survey. EClinicalMedicine, 100769 (2021).

40. J. Hadjadj et al., Impaired type I interferon activity and inflammatory responses in severe COVID-19 patients. Science 369, 718–724 (2020).

41. I. E. Galani et al., Untuned antiviral immunity in COVID-19 revealed by temporal type I/III interferon patterns and flu comparison. Nature immunology 22, 32–40 (2021).

42. C. G. K. Ziegler et al., Impaired local intrinsic immunity to SARS-CoV-2 infection in severe COVID-19. Cell 184, 4713–4733 e4722 (2021).

43. P. Bastard et al., Autoantibodies against type I IFNs in patients with life-threatening COVID-19. Science 370, (2020).

44. P. Bastard, et al., Autoantibodies neutralizing type I IFNs are present in ∼4% of uninfected individuals over 70 years old and account for ∼20% of COVID-19 deaths. Sci Immunol 6, (2021).

45. E. Y. Wang et al., Diverse functional autoantibodies in patients with COVID-19. Nature 595, 283–288 (2021).

46. J. Lopez et al., Early nasal type I IFN immunity against SARS-CoV-2 is compromised in patients with autoantibodies against type I IFNs. The Journal of experimental medicine 218, (2021).

47. P. Bastard et al., Preexisting autoantibodies to type I IFNs underlie critical COVID-19 pneumonia in patients with APS-1. The Journal of experimental medicine 218, (2021).

48. E. Pairo-Castineira et al., Genetic mechanisms of critical illness in COVID-19. Nature 591, 92–98 (2021).

49. Q. Zhang et al., Inborn errors of type I IFN immunity in patients with life-threatening COVID-19. Science 370, (2020).

50. B. Israelow et al., Mouse model of SARS-CoV-2 reveals inflammatory role of type I interferon signaling. The Journal of experimental medicine 217, (2020).

51. R. Channappanavar et al., Dysregulated Type I Interferon and Inflammatory Monocyte-Macrophage Responses Cause Lethal Pneumonia in SARS-CoV-Infected Mice. Cell host & microbe 19, 181–193 (2016).

52. J. Major et al., Type I and III interferons disrupt lung epithelial repair during recovery from viral infection. Science 369, 712–717 (2020).

53. A. Broggi et al., Type III interferons disrupt the lung epithelial barrier upon viral recognition. Science 369, 706–712 (2020).

54. C. Nair Kesavachandran, F. Haamann, A. Nienhaus, Frequency of thyroid dysfunctions during interferon alpha treatment of single and combination therapy in hepatitis C virus-infected patients: a systematic review based analysis. PLoS One 8, e55364 (2013).

55. J. C. Hall, A. Rosen, Type I interferons: crucial participants in disease amplification in autoimmunity. Nature reviews. Rheumatology 6, 40–49 (2010).

56. E. Lana-Elola et al., Genetic dissection of Down syndrome-associated congenital heart defects using a new mouse mapping panel. Elife 5, (2016).

57. H. Zhang, L. Liu, J. Tian, Molecular mechanisms of congenital heart disease in down syndrome. Genes Dis 6, 372–377 (2019).

58. C. R. Balistreri et al., Susceptibility to Heart Defects in Down Syndrome Is Associated with Single Nucleotide Polymorphisms in HAS 21 Interferon Receptor Cluster and VEGFA Genes. Genes (Basel) 11, (2020).

59. Z. Ye et al., Maternal Viral Infection and Risk of Fetal Congenital Heart Diseases: A Meta-Analysis of Observational Studies. J Am Heart Assoc 8, e011264 (2019).

60. L. J. Yockey, A. Iwasaki, Interferons and Proinflammatory Cytokines in Pregnancy and Fetal Development. Immunity 49, 397–412 (2018).

61. L. Adang et al., Developmental Outcomes of Aicardi Goutieres Syndrome. J Child Neurol 35, 7–16 (2020).

62. A. L. Rachubinski et al., Janus kinase inhibition in Down syndrome: 2 cases of therapeutic benefit for alopecia areata. JAAD Case Rep 5, 365–367 (2019).

63. B. Pinto et al., Rescuing Over-activated Microglia Restores Cognitive Performance in Juvenile Animals of the Dp(16) Mouse Model of Down Syndrome. Neuron 108, 887–904 e812 (2020).

64. F. Guedj et al., Apigenin as a Candidate Prenatal Treatment for Trisomy 21: Effects in Human Amniocytes and the Ts1Cje Mouse Model. Am J Hum Genet 107, 911–931 (2020).

65. C. L. Hunter, D. Bachman, A. C. Granholm, Minocycline prevents cholinergic loss in a mouse model of Down’s syndrome. Ann Neurol 56, 675–688 (2004).

66. L. A. Liggett et al., Precocious clonal hematopoiesis in Down syndrome is accompanied by immune dysregulation. Blood Adv 5, 1791–1796 (2021).

67. B. Bushnell, J. Rood, E. Singer, BBMerge - Accurate paired shotgun read merging via overlap. PLoS One 12, e0185056 (2017).

68. D. Kim, J. M. Paggi, C. Park, C. Bennett, S. L. Salzberg, Graph-based genome alignment and genotyping with HISAT2 and HISAT-genotype. Nature biotechnology 37, 907–915 (2019).

69. H. Li et al., The Sequence Alignment/Map format and SAMtools. Bioinformatics 25, 2078–2079 (2009).

70. S. Anders, P. T. Pyl, W. Huber, HTSeq--a Python framework to work with high-throughput sequencing data. Bioinformatics 31, 166–169 (2015).

71. A. Subramanian et al., Gene set enrichment analysis: a knowledge-based approach for interpreting genome-wide expression profiles. Proc Natl Acad Sci U S A 102, 15545–15550 (2005).

72. A. A. Sergushichev, An algorithm for fast preranked gene set enrichment analysis using cumulative statistic calculation. bioRxiv, 060012 (2016).

73. A. Liberzon et al., The Molecular Signatures Database (MSigDB) hallmark gene set collection. Cell Syst 1, 417–425 (2015).

74. K. D. Sullivan et al., The COVIDome Explorer researcher portal. Cell reports 36, 109527 (2021).

75. H. Wickham, ggplot2: Elegant graphics for data analysis (Springer -Verlag). (2016).

76. G. E. Truett et al., Preparation of PCR-quality mouse genomic DNA with hot sodium hydroxide and tris (HotSHOT). Biotechniques 29, 52, 54 (2000).

77. C. Xie, M. T. Tammi, CNV-seq, a new method to detect copy number variation using high-throughput sequencing. BMC bioinformatics 10, 80 (2009).

78. A. M. Kruisbeek, in Current Protocols in Immunology, A. M. K. J.E. Coligan, D.H. Margulies, E.M. Shevach, and W. Strober, Ed. (John Wiley & Sons, Inc., United States, 1993), chap. 3.1, pp. 3.1.1–3.1.5.

79. Z. Li et al., Duplication of the entire 22.9 Mb human chromosome 21 syntenic region on mouse chromosome 16 causes cardiovascular and gastrointestinal abnormalities. Human molecular genetics 16, 1359–1366 (2007).

80. S. S. Nandi, P. K. Mishra, Harnessing fetal and adult genetic reprograming for therapy of heart disease. J Nat Sci 1, (2015).

81. M. J. Castelhano-Carlos, N. Sousa, F. Ohl, V. Baumans, Identification methods in newborn C57BL/6 mice: a developmental and behavioural evaluation. Lab Anim-Uk 44, 88–103 (2010).

82. J. W. Goodliffe et al., Absence of prenatal forebrain defects in the Dp (16) 1Yey/+ mouse model of Down syndrome. Journal of Neuroscience 36, 2926–2944 (2016).

83. S. F. Kazim, J. Blanchard, R. Bianchi, K. Iqbal, Early neurotrophic pharmacotherapy rescues developmental delay and Alzheimer’s-like memory deficits in the Ts65Dn mouse model of Down syndrome. Scientific reports 7, 45561 (2017).

84. R. Kroeker, G. Sackett, J. Reynolds, Statistical methods for describing developmental milestones with censored data: effects of birth weight status and sex in neonatal pigtailed macaques. Am J Primatol 69, 1313–1324 (2007).

85. J. P. Lougheed, L. Benson, P. M. Cole, N. Ram, Multilevel survival analysis: Studying the timing of children’s recurring behaviors. Dev Psychol 55, 53–65 (2019).

86. P. M. G. Terry M. Therneau, Modeling Survival Data: Extending the Cox Model. . Statistics for Biology and Health (Springer, New York, NY, ed. 1, 2000), pp. XIV, 350.

87. T. M. Therneau, P. M. Grambsch, V. S. Pankratz, Penalized Survival Models and Frailty. Journal of Computational and Graphical Statistics 12, 156–175 (2003).

88. H.-P. Piepho, An Algorithm for a Letter-Based Representation of All-Pairwise Comparisons. Journal of Computational and Graphical Statistics 13, 456–466 (2004).

89. A. H. a. S. C. David Robinson. (https://CRAN.R-project.org/package=broom, 2021).

90. N. M. Aziz et al., Lifespan analysis of brain development, gene expression and behavioral phenotypes in the Ts1Cje, Ts65Dn and Dp (16) 1/Yey mouse models of Down syndrome. Disease models & mechanisms 11, dmm031013 (2018).

91. R. D’Hooge, P. P. De Deyn, Applications of the Morris water maze in the study of learning and memory. Brain Research Reviews 36, 60–90 (2001).

92. R. G. M. Morris, Spatial localization does not require the presence of local cues. Learning and Motivation 12, 239–260 (1981).

93. T. Yu et al., Effects of individual segmental trisomies of human chromosome 21 syntenic regions on hippocampal long-term potentiation and cognitive behaviors in mice. Brain Research 1366, 162–171 (2010).

94. L. Zhang et al., Human chromosome 21 orthologous region on mouse chromosome 17 is a major determinant of Down syndrome-related developmental cognitive deficits. Human Molecular Genetics 23, 578–589 (2014).

95. S. Lele, J. T. Richtsmeier, Euclidean distance matrix analysis: a coordinate-free approach for comparing biological shapes using landmark data. Am J Phys Anthropol 86, 415–427 (1991).

96. RStudio Team. (RStudio, PBC, Boston, MA, 2020).

97. R Core Team. (R Foundation for Statistical Computing, Vienna, Austria, 2020).

